# Spatial metabolic gradients in the liver and small intestine

**DOI:** 10.1101/2025.06.02.657306

**Authors:** Laith Z. Samarah, Clover Zheng, Xi Xing, Won Dong Lee, Amichay Afriat, Uthsav Chitra, Michael MacArthur, Wenyun Lu, Connor S. R. Jankowski, Cong Ma, Craig J. Hunter, Benjamin J. Raphael, Joshua D. Rabinowitz

## Abstract

Mammalian organs are composed of cells, whose properties differ depending upon their spatial location. Gene expression in liver varies between periportal and pericentral hepatocytes^1–3^, and in intestine from crypts to villus tips^4,5^. A key element of tissue spatial organization is likely metabolic, but direct assessments of spatial metabolism remain limited. Here we map spatial metabolic gradients in murine liver and intestine. We developed an integrated experimental-computational workflow using MALDI imaging mass spectrometry, isotope tracing, and deep-learning artificial intelligence. Most measured metabolites (> 95%) showed significant spatial concentration gradients in liver lobules and intestinal villi. In the liver, tricarboxylic acid (TCA)-cycle metabolites and their labeling from both glutamine and lactate localized periportally. Energy-stress metabolites including adenosine monophosphate (AMP) also localized periportally, consistent with high periportal energy demand. In intestine, the TCA intermediates malate (tip) and citrate (crypt) showed opposite spatial patterns, which aligned with higher glutamine catabolism in tips and lactate oxidation in crypts based on isotope tracing. Finally, we mapped the fate of the obesogenic dietary sugar fructose. In the intestine, oral fructose was catabolized faster in the villus bottom than the tips. In the liver, fructose-derived carbon accumulated pericentrally as fructose-1-phosphate and triggered pericentral adenosine triphosphate (ATP) depletion. Thus, we both provide foundational knowledge regarding intestine and liver metabolic organization and identify fructose-induced focal derangements in liver metabolism.

## Introduction

In mammals, tissue metabolism is fueled by circulating nutrients that are supplied directly from the diet or through inter-organ exchange^6,7^. The two visceral organs, small intestine and liver, occupy strategically important sites. Located at the front line of the digestive system, the small intestine contributes to digestion and nutrient absorption^8^, while shielding other organs by catabolizing dietary nutrients^9^. The liver, in addition to processing dietary nutrients and xenobiotics, is a key regulator of carbohydrate, fat, and protein metabolism^10–12^. Together, both organs orchestrate consumption, production, and distribution of dietary and circulating nutrients, thereby shaping whole-body metabolic homeostasis.

Tissue function is strongly influenced by its spatial architecture. For example, in the liver, cells are spatially organized into repeating hexagonal units called lobules^13,14^. Portal triads (equivalently, ‘portal nodes’) are located at the lobule corners and consist of a portal vein, hepatic artery, and bile duct. Nutrient-rich and oxygenated blood enters each lobule through the hepatic artery and portal vein and drains in the central vein located at the lobular center. Such spatial arrangement induces a concentration gradient of nutrients and oxygen along the portal-central axis. Hepatocyte gene expression similarly varies along the portal-central axis^2,3,10,15^, supporting zonation of the liver into periportal and pericentral regions. Such spatial zonation helps the liver in performing a wide range of physiological processes simultaneously, some of which are functionally reciprocal^1,3,10^. For example, the liver both catabolizes and synthesizes glutamine. Based on gene expression, glutamine breakdown is periportal, whereas resynthesis is confined to hepatocytes contiguous to the central vein^1,2^.

Spatial gene-expression gradients are also observed in the small intestine^5,16,17^. There, differentiated epithelial cells originate in the crypts and migrate towards the villus tips^18–20^. Cells along the crypt-villus axis are exposed to concentration gradients of circulating compounds as blood flows from the submucosal layer to the villus tip. Analogous to hepatocytes along the portal-central axis, enterocytes display spatial gene-expression gradients along the crypt-villus axis^5,16^.

Up to now, metabolic processes in the liver and small intestine have been spatially inferred largely from gene and protein expression^1,2,5,21,22^. Around 50% and 80% of liver and small-intestine genes, respectively, have been found to be zonated^2,5^. Gene expression, however, does not always reflect cellular phenotype. Protein and transcript levels are only weakly spatially correlated^21,22^. Moreover, enzyme levels only partially explain metabolic activity^23^. A few studies have explored spatial distributions of metabolites^24^ and lipids^25^ in the liver using imaging mass spectrometry (IMS), while measurements of spatial metabolic gradients in the small intestine are yet to be achieved. To date, liver spatial metabolomics has largely been confined to low spatial resolution (sampling area ≥ 30 × 30 µm^2^) and has not quantified porto-central gradients^24^, with the exception of a few metabolites measured through secondary ion mass spectrometry, a technique with micron spatial resolution but limited metabolome coverage^26^. Furthermore, quantifying spatial gradients holds potential for assessing local metabolic derangements associated with physiological perturbations.

Beyond metabolite abundances, there is also great potential in understanding metabolic activity spatially. Metabolite levels alone do not reflect pathway activity^27–31^, as higher metabolite abundances can be attributed to either faster production or slower consumption. The combination of stable-isotope tracing and imaging mass spectrometry hold potential in this regard^32–34^, but thus far has only been achieved at low spatial resolution for tracing core small-molecule metabolites.

Here, we map spatial metabolic gradients in murine liver and intestine. We measure metabolite abundances and pathway activity at high spatial resolution (15 µm for liver; 10 µm for intestine) using matrix-assisted laser desorption ionization - imaging mass spectrometry (MALDI-IMS). We develop a deep-learning approach that infers from the metabolomic images in an unbiased manner the predominant underlying spatial pattern (metabolic topography). In the liver, this recapitulates classical portal-central organization and, in intestine, crypt-villus organization. More than 95% of metabolites vary significantly along these classical tissue axes, establishing metabolic gradients as a hallmark of tissue organization. We further show metabolic activity patterns underlying these concentration gradients, and local disruption in the levels of perhaps the most fundamental metabolite, ATP, by the obesogenic dietary nutrient fructose.

## Results

### Deep learning of liver metabolic topography

We collected MALDI-IMS data from slices of flash-frozen murine liver. Data were collected in negative ion mode at 15-µm spatial resolution using N-(1-Naphthyl) ethylenediamine dihydrochloride (NEDC) as the matrix^35^. From the raw data, around 170 ions, including those corresponding to deprotonated and other forms of canonical lipids and water-soluble metabolites, were reliably detected. Of these, around 120 were deprotonated ions of canonical metabolites based on exact mass match and further identity confirmation (natural isotope abundance distribution, tandem mass spectrum, and/or isotope labeling; Supplementary Table 1).

A common challenge in IMS is to assign pixels a histological identity, which can sometimes be tackled by co-registering and overlaying spatial information from other imaging modalities^36–39^. In the context of liver anatomy, we sought to determine the spatial location of a given IMS pixel along the liver portal-central axis in an automated manner and without the need for other orthogonal imaging approaches (Fig. 1a). Both portal and central veins could be identified by the presence of heme (Fig. 1c). Portal nodes were further distinguished by the presence of bile ducts as indicated by bile-acid ions (taurocholic acid) (Fig. 1c) and confirmed by histology of the same slide post-MALDI imaging (Fig. 1b) and by immunofluorescence imaging (Extended Data Fig. 1a).

**Figure 1.**
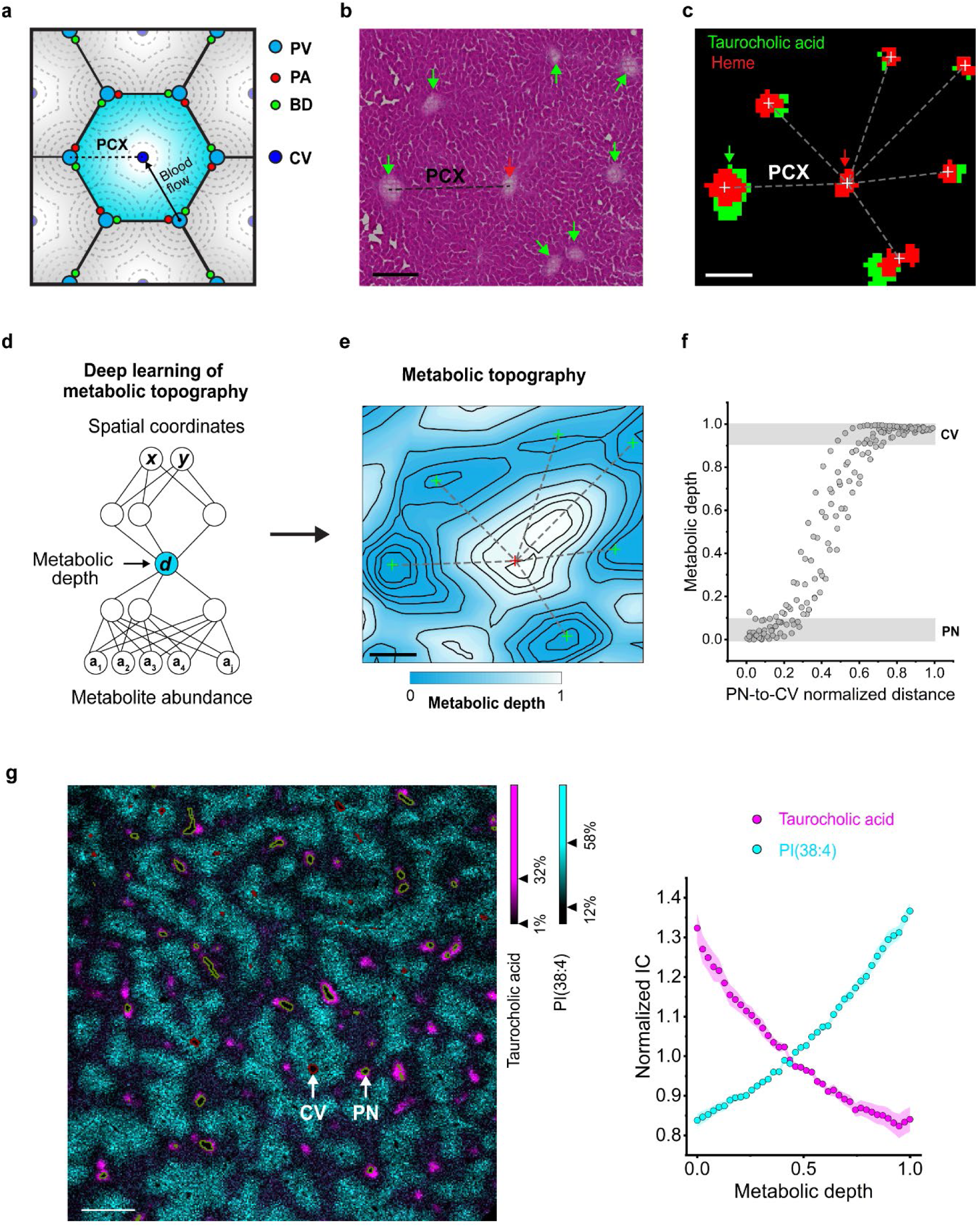
Deep-learning neural network for inferring liver metabolic topography. **(a)** Illustration of a liver lobule (blue hexagon). PV, portal vein; PA, portal artery; BD, bile duct; CV, central vein; PCX, portal-central axis. **(b)** H&E of liver tissue slice post MALDI-IMS. Red arrow, central vein; green arrows, portal nodes (PN); dashed line, portal-central axis (PCX). **(c)** Merged binarized MALDI images of heme (red) and taurocholic acid (green) in liver from (b). Red and green arrows represent central vein and portal node, respectively. Crosses represent centroids of assigned veins and dashed lines PCX. **(d)** Neural network learns the one-dimensional coordinate ‘metabolic depth’ associated with each spatial location based on ion counts of reliably detected metabolite ions. **(e)** Metabolic topography shown as a contour map with contour lines of constant metabolic depth. Vein positions from (c) are overlaid. **(f)** Correlation between metabolic depth and geometrical normalized distance from portal node to central vein. Data are for pixels located along the portal-central axes in (e). Gray bands represent metabolic depths associated with blood vessels rather than liver parenchyma and are omitted from subsequent analyses of metabolic gradients. **(g)** Overlaid images of ion counts for taurocholic acid (magenta) and PI(38:4) (cyan) (intravascular areas, defined by high heme, are excluded). Outlines of portal veins are olive green and central veins are wine red. PN, portal node. Right panel: Ion counts (IC) as a function of metabolic depth. Pixels were divided by metabolic depth into 50 bins. Each dot is the mean ion count of pixels in that metabolic depth bin across n = 7 independent mice and error band is the standard error of the mean (SEM). Scale bars (b – e) = 150 µm, (g) = 600 µm. MALDI images were collected at 15 × 15 µm^2^ spatial resolution.

We hypothesized that position along the portal-central axis would be the primary spatial metabolic feature in liver, and that it could be automatically inferred by deep learning. Building from our recently introduced method GASTON for spatial transcriptomics^40^, we developed a deep-learning method for identifying spatial metabolic gradients (Metabolic Topography Mapper, MET-MAP) (Fig. 1d). This approach learns a one-dimensional coordinate (‘metabolic depth’) in an unsupervised manner (i.e., without prior knowledge of tissue anatomy) that best recapitulates the observed high-dimensional spatial metabolomics data (Fig. 1d,e). Metabolic depth is analogous to contour lines on a topographical map. In liver, it recapitulates the classical hexagonal lobule architecture (Fig. 1e) and correlates strongly with position measured directly along the observable portal-central axes (Fig. 1f and Extended Data Fig. 1b-e). Thus, in a fully unsupervised manner, metabolic depth defined the hallmark portal-central organization of the liver.

We next examined how metabolite concentrations vary as a function of metabolic depth. Taurocholic acid and PI(38:4) (phosphatidylinositol with 38 fatty acyl carbons and 4 degrees of unsaturation) are representative metabolites showing classical portal-central gradients (Fig. 1g). Regression, omitting extreme values of metabolic depth to focus on hepatocytes and exclude veins, then provides a convenient way to identify significant portal-central gradients. Negative slopes reflect periportal localization (as for taurocholic acid), and positive ones pericentral localization (Fig. 1g). Thus, we established a deep-learning method for inferring liver metabolic topography that identifies metabolites with periportal or pericentral localization.

### Periportal localization of TCA and energy-stress metabolites

We next assessed which metabolites showed significant portal-central gradients. To this end, metabolite signal intensity was plotted versus metabolic depth, using MALDI-IMS data collected from 7 independent mice. Of the total 120 metabolites and lipids, over 95% exhibited statistically significant spatial gradients (*p* < 0.01 for slopes from linear regression with Benjamini-Hochberg FDR correction) (Fig. 2a and Supplementary Table 2). Many fatty acids (which may come from in-source fragmentation of lipids) and phospholipids were pericentrally localized (Fig. 2a).

**Figure 2.**
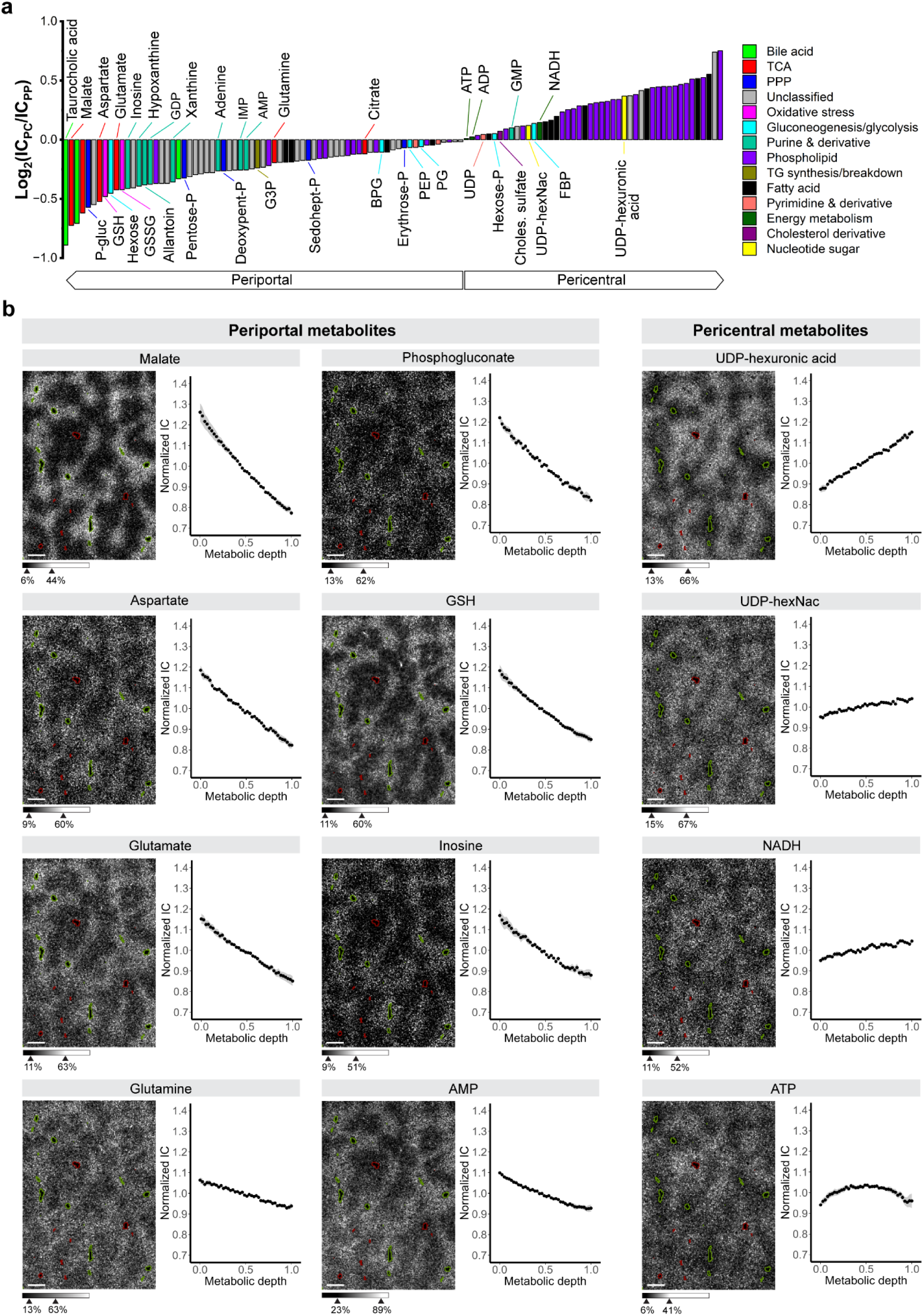
Liver portal-central metabolite concentration gradients. **(a)** Metabolite concentration gradients represented as pericentral (PC) to periportal (PP) fold change of ion count (log_2_ scale). Negative log_2_ fold change = periportally enriched; positive = pericentral. Pixel positions are based on metabolic depth learned from the deep neural network as in Fig. 1. Data from n = 7 independent livers were analyzed by linear regression to look for portal-central concentration trend. For all metabolites shown, *p*-value from regression < 0.01 after Benjamini-Hochberg FDR correction. For individual *p*-values, see Supplementary Table 2. **(b)** Individual metabolite raw ion count images and their respective spatial gradients (graphs to right of images). In the graphs, data points reflect the mean and shading SEM across n = 7 independent mice. Images are as in Fig. 1g (green outline = portal vein, red = central vein), with ion intensities in gray scale. Scale bar = 300 µm. MALDI images were collected at 15 × 15 µm^2^ spatial resolution. Compound name abbreviations: P-gluc, phosphogluconate; Pentose-P, pentose phosphate; Deoxypent-P, deoxypentose phosphate; Sedohept-P, sedoheptulose phosphate; Erythrose-P, erythrose phosphate; BPG, bisphosphoglycerate; PEP, phosphoenolpyruvate; PG, phosphoglycerate; G3P, glycerol 3-phosphate; Hexose-P, hexose phosphate; Cholest. sulfate, cholesterol sulfate; UDPhexNAc, UDP-N-acetylhexosamine; FBP, fructose bisphosphate; GSH, reduced glutathione; GSSG, oxidized glutathione.

Glucose is both produced and consumed by the liver. Based on gene expression, gluconeogenesis tends to be periportal and glycolysis pericentral^1,2,10^. Consistent with glucose being made periportally and consumed pericentrally, its levels were higher periportally (Fig. 2a and Extended Data Fig. 2a), while the downstream glycolytic intermediates glucose-6-phosphate (together with its isomers, ‘hexose phosphate’) and fructose bisphosphate both localized pericentrally (Fig. 2a).

One important use of glucose-6-phosphate is synthesis of UDP-sugars. UDP-glucose is the direct substrate for glycogen synthesis and precursor of the other UDP-sugars, two of which were detectable by MALDI imaging: UDP-glucuronic acid, which is used for xenobiotic glucuronidation and thus detoxification^41,42^, and UDP-N-acetylglucosamine, for protein glycosylation^43,44^. These localized pericentrally (Fig. 2a,b), as did, based on published spatial transcriptomics and proteomics data, the enzymes driving these processes^2^ (Extended Data Fig. 3a). Thus, substrates and enzymes for both protein glycosylation and xenobiotic detoxification colocalize pericentrally.

Another important consumer of glucose-6-phosphate is the pentose phosphate pathway (PPP), which makes NADPH and the nucleotide precursor ribose-5-phosphate (together with its isomers, ‘pentose phosphate’). The PPP intermediates 6-phosphogluconate, pentose phosphate, and sedoheptulose-7-phosphate all localized periportally, as did PPP enzymes (Fig. 2a,b and Extended Data Fig. 3b). A major function of NADPH is maintenance of glutathione in its reduced state, and reduced glutathione (GSH) was also periportal (Fig. 2a,b and Extended Data Fig. 2b).

Although glycolysis generates some energy, most ATP in mammals is made oxidatively, through the tricarboxylic acid (TCA) cycle coupled to the electron transport chain^45^. In the liver, oxygen levels drop as blood flows from the portal triad to the central vein^46,47^. This reflects oxidative periportal metabolism, with both TCA-cycle enzymes (Extended Data Fig. 3c) and mitochondrial density^48,49^ previously shown to be greater periportally. Consistent with this, TCA- cycle intermediates localized periportally, as did the amino acids aspartate and glutamate which are in rapid exchange with TCA intermediates (Fig. 2a,b). This periportal localization was strongest for the four-carbon metabolites malate and aspartate (Fig. 2a,b). Gluconeogenesis involves flux through four-carbon TCA intermediates but not the rest of the TCA cycle; thus, the strong localization of these metabolites is consistent with periportal gluconeogenesis. A key output of the TCA cycle is NADH, which notably localized pericentrally, opposite to TCA intermediates, perhaps due to lower oxygen and thus slower oxidative phosphorylation in the pericentral region (Fig. 2a,b).

TCA cycle and oxidative phosphorylation operate to meet energy demand, which is high periportally due to urea cycle and gluconeogenesis. Intuitively, one might expect high periportal TCA activity to result in high ATP and energy charge (approximated here by the ATP/AMP ratio). Our spatial images reveal the opposite. ATP is higher pericentrally and AMP periportally (Fig. 2a,b and Extended Data Fig. 2c). Despite some AMP signal in MALDI arising from ATP fragmentation (and thus AMP signal alone having multiple interpretations), the combination of lower ATP and higher AMP periportally reliably indicates low energy charge. When AMP rises, it is catabolized by AMP deaminase in an effort to maintain energy charge, producing inosine monophosphate (IMP), inosine, hypoxanthine, and xanthine. All of these catabolites localized periportally (Fig. 2a,b). Thus, despite high periportal levels of mitochondria and TCA intermediates, periportal energy demand appears to be sufficient to trigger energy stress.

### Metabolic topography of the intestinal epithelium

Similar to repeating lobules in the liver, the small intestine consists of recurring crypt-villus units. Each villus is covered in epithelial cells, mainly enterocytes, surrounding blood vessels, lymphatics, and lamina propria^20^ (Fig. 3a). Enterocytes are exposed to oxygen and nutrient gradients along the crypt-villus axis, with oxygen enriched in the crypts and consumed as blood flows towards the tips. We sought to quantify spatial metabolic gradients in the epithelial-cell layer along the crypt-villus axis. Substantial methodological advancement was required to capture adequate MALDI images for this purpose. The epithelium is only a single-cell layer, so high spatial resolution is required. Moreover, in contrast to liver slices, the intestine is hollow, requiring tissue embedding. To facilitate single-cell resolution MALDI imaging, the resulting slices were coated with the MALDI matrix 1, 5-diaminonaphthalene (DAN), which has small and homogeneous matrix crystal size, as required for obtaining high spatial resolution MALDI images^50^. The resulting slides were analyzed by MALDI-IMS at 10-µm spatial resolution with around 100 metabolite ions reliably detected (Supplementary Table 3).

**Figure 3.**
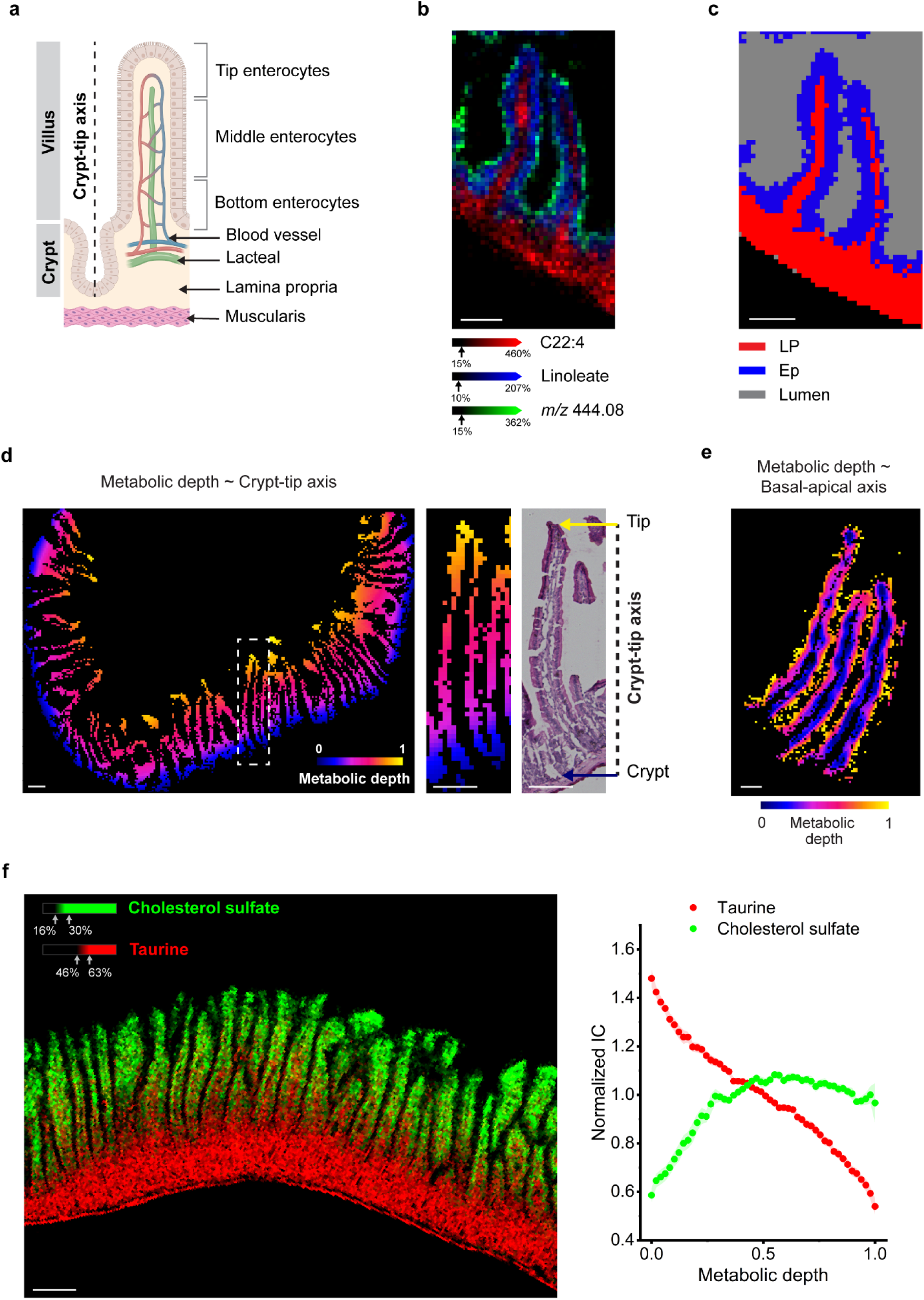
High resolution metabolic imaging of small intestine epithelium. **(a)** Illustration of small-intestine anatomy. **(b)** The fatty acid C22:4 (red) highlights the lamina propria, linoleate (C18:2, blue) represents the epithelial layer, and ion at *m/z* 444.08 (putatively annotated as a mucin fragment^73^) localizes in the mucin layer covering the epithelium. **(c)** Anatomical regions are segmented by k-means clustering using data from about 100 metabolites that are reliably detected in intestine. Clustering captures the lamina propria (LP; red), epithelial layer (Ep; blue) and lumen (gray). **(d)** Epithelial-layer data is used by the deep neural network to learn two metabolic depths. The first metabolic depth is shown in (d) and captures villus crypt-to-tip metabolite variation, validated by H&E staining (right panel). Dotted rectangle in the left panel highlights the villus magnified in the center panel. **(e)** The second metabolic depth captures the epithelial cell brush broader-to-basal membrane (villus exterior-to-interior) metabolite gradient. **(f)** Overlaid images of ion counts for taurine (red) and cholesterol sulfate (green) and their corresponding ion-count variation as a function of the first metabolic depth. Data shown in graphs was processed as in Figure 2 and is the mean and SEM for n = 7 mice (see Methods for details). (b – d) Scale bar = 100 µm; (e) 120 µm; (f) 500 µm. All MALDI images shown were collected at 10 × 10 µm^2^ spatial resolution, except for (e), which was acquired at 6 × 6 µm^2^ spatial resolution.

We sought to implement our deep-learning approach to quantify metabolic gradients in epithelial cells residing along the crypt-villus axis. Unlike in liver tissue slices, where hepatocytes predominate, intestine cross sections contain mainly lumen and lamina propria, with the epithelial cells only a thin layer. Bile acids are secreted by the liver into the intestinal lumen to facilitate dietary fat absorption. We observed high bile acid (e.g., taurocholic acid) signal in the lumen and fatty acid signals in the intestinal epithelium and lamina propria (Fig. 3b and Extended Data Fig. 4a). Specific fatty acids distinguished intestinal epithelium (linoleate, C18:2) from lamina propria (C22:4) (Fig. 3b and Extended Data Fig. 4a,b). Motivated by these patterns, we used k-means clustering to group all pixels into clusters based on the roughly 100 quantifiable ions, (Fig. 3c). This yielded reliable categorization of pixels as background, lumen, lamina propria, and epithelium, confirmed by H&E-staining post-MALDI imaging (Extended Data Fig. 4b).

Visual inspection of the epithelial-cell IMS data suggested two potential types of gradients: villus crypt-to-tip and epithelial cell brush border-to-basal membrane. We accordingly developed a version of MET-MAP capable of assessing two different metabolic gradients simultaneously (the output for each pixel being two different metabolic depth values) (Fig. 3d,e). In all intestine metabolomic images, one metabolic depth corresponded to the crypt-villus axis (Fig. 3d) and was used for subsequent analyses. To identify metabolites with significant crypt-villus gradients, we examined ion count as a function of crypt-villus metabolic depth, with positive slopes indicating tip enrichment. Taurine, one of the most abundant small-molecule metabolites in mammals, localized strongly to crypts (Fig. 3f), where it may help protect against oxidative stress^51^. In contrast, cholesterol sulfate localized to tips (Fig. 3f), where it may contribute to stabilizing the membranes of aging enterocytes^52^.

### Spatial discordance in TCA and energy metabolites along crypt-villus axis

Overall, around 95% of detected metabolites exhibited statistically significant spatial gradients in intestinal epithelium from crypt-to-tip (*p* value < 0.01 by linear regression with Benjamini-Hochberg FDR correction) (Fig. 4a and Supplementary Table 4). These included 55 fatty acid and lipid species, with the majority localizing in crypts and villus bottom (Fig. 4a). Overall, degree of unsaturation in fatty acyls was greatest at the crypts (Fig. 4a). For example, C20:3 strongly marked the villus bottom (Fig. 4b). This correlation may reflect loss of highly unsaturated tails to oxidative damage as enterocytes migrate from crypt to tip.

**Figure 4.**
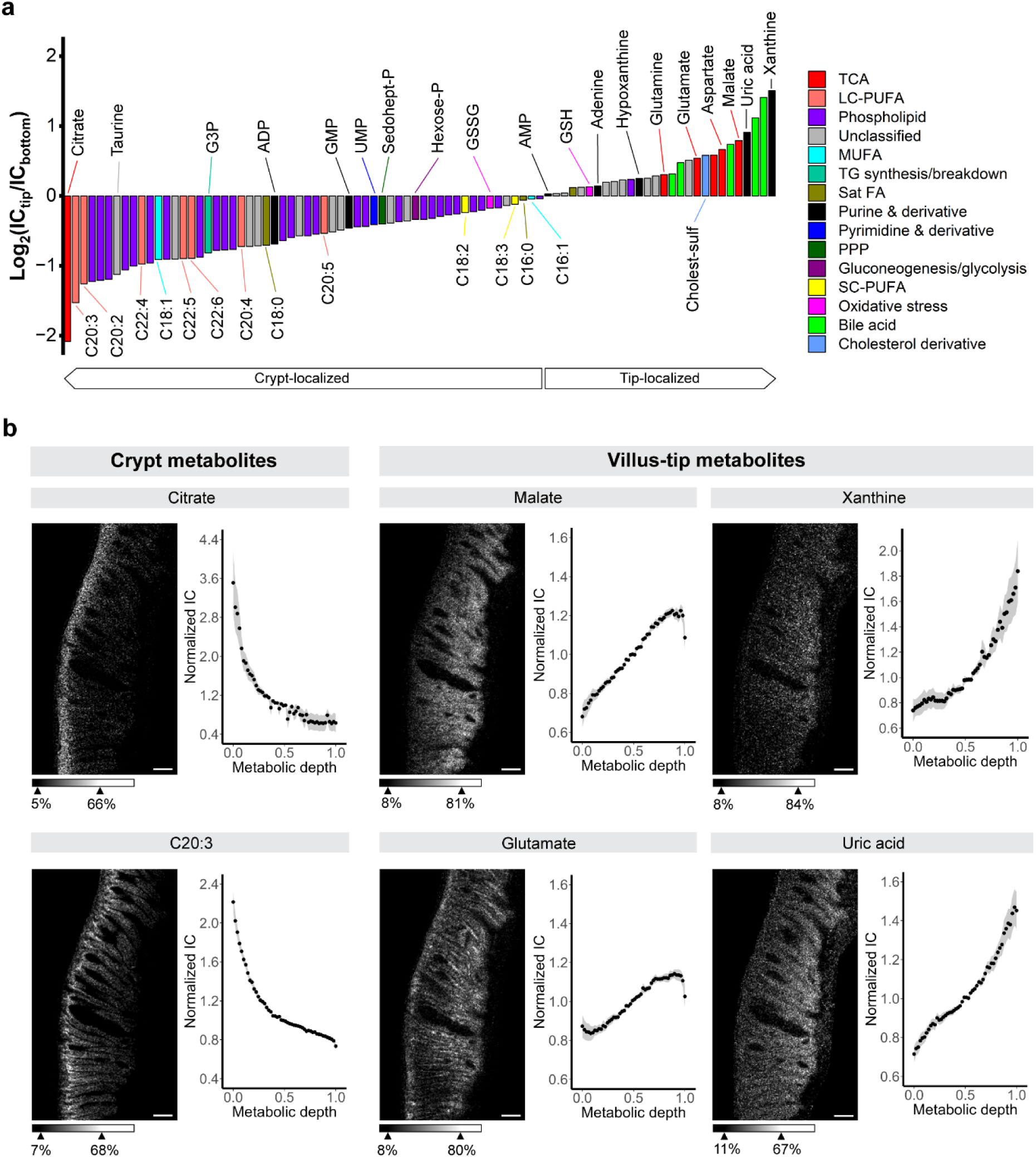
Opposing spatial patterns of TCA intermediates in intestinal epithelium. **(a)** Metabolite concentration gradients represented as crypt-to-tip fold change of ion count (log_2_ scale). Negative log_2_ fold change = crypt-enriched; positive = tip. Data from n = 7 independent intestines and analyzed by linear regression to look for crypt-tip concentration trend. For all metabolites shown, *p*-value from regression < 0.01 after Benjamini-Hochberg FDR correction; for individual *p*-values, see Supplementary Table 3. **(b)** Individual metabolite raw ion count images and their respective spatial gradients (graphs to right of images). Data points reflect the mean and shading SEM across n = 7 independent mice. Scale bar = 200 µm. MALDI images were collected at 10 × 10 µm^2^ spatial resolution. For compound name abbreviations, see Fig. 2.

In terms of water-soluble metabolites, the most strongly localized were TCA intermediates (Fig. 4a,b). Citrate localized to crypts (Fig. 4a,b). In contrast, malate localized to tips (Fig. 4a,b), as did the amino acids aspartate and glutamate (Fig. 4a,b), which are in rapid exchange with TCA intermediates. Mitochondria and their genes tend to localize to tips^19^ (Extended Data Fig. 6a). As in the liver, this colocalizes malate, aspartate and glutamate with mitochondria. The opposite spatial localization of citrate is not readily explained by enzyme expression patterns, as citrate synthase is tip localized^5^ (Extended Data Fig. 6a), consistent with its mitochondrial localization. Moreover, one of the most strongly crypt-localized enzymes is citrate lyase, which consumes citrate^5,22^ (Extended Data Fig. 6a). Thus, in the intestine, TCA metabolites show a spatially bifurcated pattern with citrate and malate localizing in opposite ends of villi.

AMP and ADP also showed opposite spatial patterns in the intestine (Fig. 4a). ADP localized in crypts where AMP was depleted. When AMP rises, it is degraded into hypoxanthine (Fig. 4a), xanthine (Fig. 4a,b), and uric acid (Fig. 4a,b), all of which were substantially tip localized. Collectively, these data suggest lower energy charge in the tips. The key energy carrier ATP itself was not detectable in intestine in our primary MALDI analysis but could be measured after washing the slides with organic solvent (Extended Data Fig. 4c)^53^, which enhances signals for phosphorylated compounds. ATP colocalized with ADP towards the crypts, confirming an elevated AMP/ATP ratio (i.e. low energy charge) at the villus tips (Extended Data Fig. 4c). Thus, perhaps due to high energy demand for nutrient uptake or impaired energy production due to oxygen limitation, villus tips experience energy stress.

### Substrates driving spatial TCA metabolic gradients

The above analysis identified spatial gradients in the concentrations of most measured metabolites in both liver and intestine, with particularly strong gradients in TCA-related metabolites (Fig. 2a,b and Fig. 4a,b). To investigate the underlying metabolic fluxes, we turned to isotope tracing. We systemically infused [U-^13^C,^15^N]glutamine and [U-^13^C]lactate separately to probe their pseudo-steady-state contributions to TCA-related compounds (Extended Data Fig. 5a,d). These experiments require measuring isotope-labeled forms of metabolites that are substantially less abundant than the parent compounds. In the liver, adequate data quality was obtained for both malate and glutamate, and in the intestine only for glutamate.

Among the strongest spatial enzyme gradients in murine liver is periportal localization of the glutamine catabolic enzyme glutaminase (Extended Data Fig. 5b,c). Consistent with such localization, we observed increased labeling of both glutamate (Fig. 5a,b) and malate (Fig. 5a,c) from glutamine periportally. Labeling from lactate was more modestly periportal (Fig. 5d,e,f). Lactate enters the TCA cycle via pyruvate through two different routes. Oxidative catabolism to acetyl-CoA via pyruvate dehydrogenase generates M+2 TCA intermediates and supports TCA turning and energy generation (Fig. 5d). In contrast, pyruvate carboxylation to oxaloacetate generates M+3 TCA intermediates and feeds gluconeogenesis (Fig. 5d). Consistent with higher gluconeogenic flux periportally, malate M+3/M+2 ratio was greater periportally (Fig. 5f). We were further able to track resynthesis of glutamine from glutamate, which is catalyzed by the strongly pericentrally localized enzyme glutamine synthetase (Extended Data Fig. 5c), finding that this metabolic flux localizes pericentrally (Fig. 5a,b and Extended Data Fig. 5c). Overall, these tracing data confirm expected periportal localization both of gluconeogenesis and glutamine catabolism, as well as the expected pericentral localization of glutamine resynthesis from glutamate.

**Figure 5.**
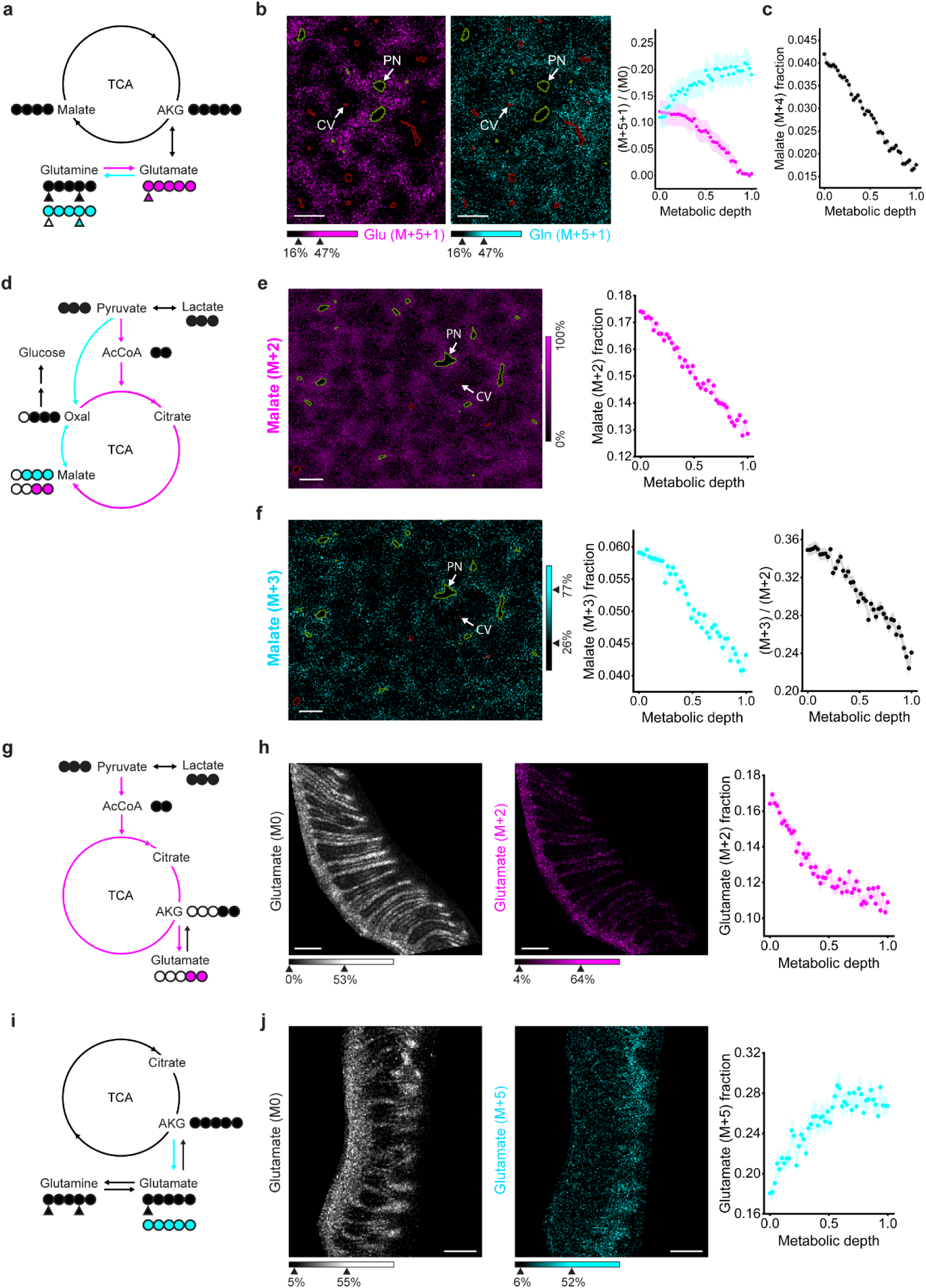
Spatial variation in TCA-cycle inputs revealed by isotope-tracer infusions. **(a)** Schematic of fully labeled glutamine ([^13^C_5_,^15^N_2_] = M+5+2) metabolism in liver. M+5+2 glutamine can be metabolized to M+5+1 glutamate (magenta) and further to M+5 α-ketoglutarate and M+4 malate (black). M+5+1 glutamate can then be recycled via glutamine synthetase to M+5+1 glutamine (cyan). Circles: carbon atoms. Triangles: nitrogen atoms. Filled (black and colored): Isotope-labeled. **(b)** Liver MALDI images based on tracing in (a) (20 × 20 µm^2^ spatial resolution). M+5+1 glutamate = magenta. M+5+1 glutamine = cyan. Jugular-vein catheterized mice were infused with fully labeled glutamine for 2.5 h (circulating tracer labeling of ∼40%). In all panels, plotted values are raw isotope ratios without correction for circulating tracer fractional enrichment (mean ± SEM, n = 6 mice). **(c)** Same for M+4 malate fraction. **(d)** As in (a), for fully labeled lactate as the tracer ([^13^C_3_] = M+3). Lactate oxidation generates M+2 malate. Anaplerosis (pyruvate carboxylation) produces M+3 malate. **(e)** MALDI liver image of M+2 malate. Mice were infused [^13^C_3_]lactate for 2.5 h (∼40% serum enrichment) (mean ± SEM, n = 4 mice). **(f)** Same for M+3 malate and M+3/M+2 malate ratio, reflecting anaplerosis relative to TCA turning. **(g)** As in (d), for intestine (10 × 10 µm^2^ spatial resolution). Due to the higher spatial resolution, only glutamate shows sufficient signal to reliably quantify its isotope labeling and is used as the TCA readout (glutamate and the TCA intermediate α-ketoglutarate are in rapid exchange). **(h)** MALDI intestine images of unlabeled (M0) and M+2 glutamate (mean ± SEM, n = 3 mice). **(i)** As in (g), for fully labeled glutamine as the tracer. **(j)** MALDI intestine images of unlabeled glutamate (M0) and M+5 glutamate from a mouse infused with fully labeled glutamine as in (b) (mean ± SEM, n = 2 mice).

In the intestine, we were particularly interested in whether isotope tracing could shed light on the differential localization of malate (tips) and citrate (crypts). Citrate synthesis from malate involves two enzymatic reactions: malate oxidation to oxaloacetate followed by addition of a two-carbon unit from acetyl-CoA, typically coming from glucose, lactate or fat oxidation (Fig. 5g). As both of these steps from malate to citrate depend on oxidative capacity, they logically might be hindered in the less well-perfused villus tips, decreasing the citrate/malate ratio in the tips. If this were the case, we would expect the TCA cycle to be fed via oxidative lactate metabolism (M+2 TCA intermediates from lactate) preferentially in the crypts and for the major alternative fuel, glutamine, to predominate at the tips (Fig. 5g,i). Indeed, we observe preferential lactate contribution to intestinal glutamate in the crypts and glutamine contribution in the villus tips (Fig. 5h,j). Thus, TCA fuel utilization varies spatially in both the liver and intestine and helps to explain observed metabolite concentration gradients.

### Fast fructose usage in the villus bottom

Up to this point, we have spatially mapped fluxes pertaining to core metabolic pathways related to lactate and glutamine, major circulating fuels. Ultimately, many circulating metabolites are derived from diet, and the small intestine and liver are at the front lines of dietary nutrient processing. Over the last century, in the United States, fructose has shifted from being a minor nutrient to major caloric source^54^. Excessive fructose consumption is associated with obesity, diabetes, hyperlipidemia, and non-alcoholic fatty liver disease/steatohepatitis^55–59^. Accordingly, understanding spatial aspects of fructose handling is medically relevant. To this end, we administered mice with [U-^13^C]fructose and [^2^H_12_]glucose in a 1:1 ratio by oral gavage at a relatively high dose of 1 g/kg body weight (for each hexose; roughly equivalent to 1L of soda in humans, normalized on a daily calories basis) (Fig. 6a). The coadministration of fructose and glucose mimics the typical human context (e.g. sucrose or high fructose corn syrup, which are both a mix of fructose and glucose). The distinct isotope labeling of fructose and glucose allows us to distinguish downstream metabolites made from these two isomeric sugars. The committed step in fructose usage is its phosphorylation by ketohexokinase (KHK) to fructose-1-phosphate (F1P) (Fig. 6a). We measured the spatial distribution of [U-^13^C]hexose phosphate (reflecting F1P) in the intestine at 90 seconds and 10 minutes post gavage. Strikingly, F1P signal appears initially in the villus bottom, with the spatial gradient less intense at 10 minutes (Fig. 6b). A simple interpretation is that fructose catabolic flux is preferentially at the villus bottom, as reflected in the intense initial gradient and emphasizing the value of pre-steady-state measurements for capturing absolute flux^60^.

**Figure 6.**
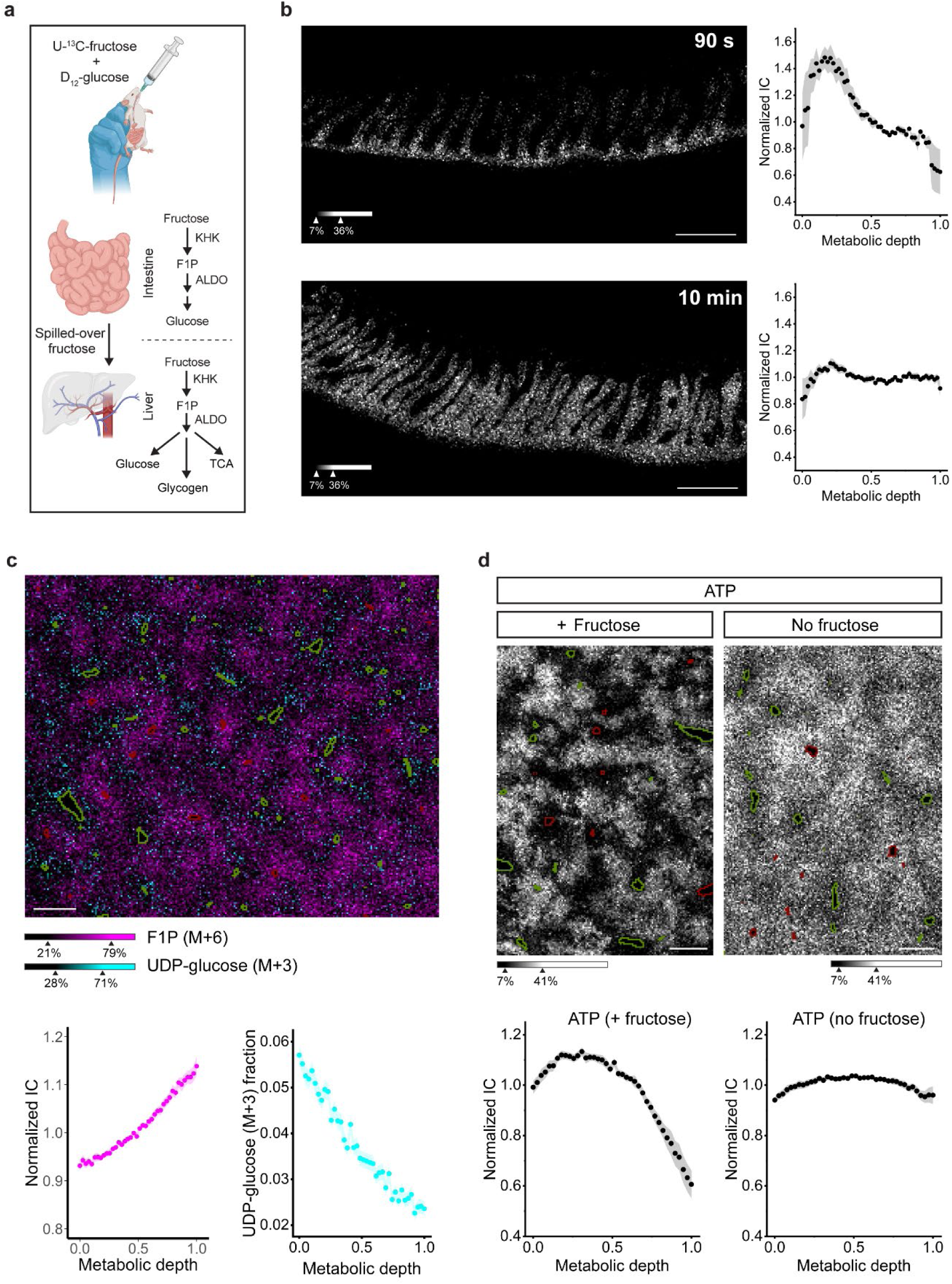
Mapping the fate of oral fructose. **(a)** Schematic of dietary fructose utilization in the small intestine and liver. The small intestine converts fructose to glucose. Fructose that is not processed by the intestine spills over to the liver through portal blood. **(b)** MALDI intestine image of M+6 fructose-1-phosphate (F1P) in the small intestine after gavage of [U-^13^C]fructose + [D_12_]glucose (1000 mg/kg each), from intestines collected post gavage at 90 seconds (top, mean ± SEM, n = 5 mice) or 10 minutes (bottom, mean ± SEM, n = 8 mice). **(c)** Overlaid MALDI liver images of M+6 F1P and M+3 UDP-glucose, 10 minutes after gavage (mean ± SEM, n = 6 mice for F1P and n = 4 mice for UDP-glucose). **(d)** MALDI liver images of unlabeled ATP, 10 minutes after fructose + glucose gavage as above, compared to untreated mice (mean ± SEM, n = 6 treated and n = 7 untreated mice). Scale bar (b) = 200 µm; (c) = 500 µm; (d) = 300 µm. MALDI images were collected at 10 × 10 µm^2^ for intestine and 20 × 20 µm^2^ or 15 × 15 µm^2^ for liver (c, d, respectively).

### Periportal glycogenesis from fructose and pericentral ATP depletion

Oral fructose that is not captured by the intestine spills over to the liver through portal blood, together with glucose and carboxylic acids generated by the intestine from oral fructose^9,61^ (Fig. 6a). The metabolites enter the liver periportally, with both the liver fructose transporter GLUT2 and KHK weakly periportal^21^ (Extended Data Fig. 5i). Unexpectedly, however, F1P (represented by [U-^13^C] hexose phosphate) was higher pericentrally (Fig. 6c). While this could reflect discordance between enzyme levels and flux, another possibility is that F1P accumulates pericentrally due to slower downstream reactions.

F1P is cleaved in the liver by aldolase B^62^, which is strongly periportal^21^ (Extended data Fig. 5i). F1P cleavage generates triose phosphates (dihydroxyacetone phosphate) that can be used either for energy production via glycolysis or sugar production/storage through gluconeogenesis. Glucose 6-phosphate made by gluconeogenesis can either be dephosphorylated for glucose release to the circulation or stored as glycogen via the key intermediate UDP-glucose. Consistent with enhanced F1P cleavage and usage periportally, we observed periportal localization of M+3 UDP-glucose, the isotopic form produced from fructose via the above series of reactions (Fig. 6c). In contrast, M+7 UDP-glucose, the form produced from co-administered 2H-glucose, tends to be pericentral, indicating that the enhanced periportal M+3 UDP-glucose is due to spatial localization of fructolysis (Extended Data Fig. 7).

One way that fructose is thought to cause metabolic damage is through ATP consumption by fructose phosphorylation, which lacks the homeostatic feedback of glycolysis^63–65^. When reactions downstream of F1P are slow and accordingly fail to effectively regenerate ATP, this can lead to energy stress. The accumulation of F1P pericentrally, where the fructolysis enzyme aldolase B is low (Extended Data Fig. 5i), suggested the possibility of imbalanced periportal ATP utilization triggered by fructose. Indeed, livers from mice that were given an acute oral dose of fructose showed striking pericentral ATP depletion (Fig. 6d). Thus, the combination of isotope tracing and high-resolution MALDI imaging of liver metabolic topography identifies fructose-induced focal metabolic derangements.

## Discussion

To date, spatial understanding of liver and intestine metabolism has been largely inferred from transcriptomics and enzyme measurements. Enzyme expression patterns, however, are insufficient to reliably predict metabolite concentrations or fluxes, reflecting the importance of other variables including substrate supply and enzyme regulation. Here, we developed integrative experimental-computational methods that enable direct assessment of tissue metabolite concentration gradients. We find that the vast majority of measured metabolites display significant spatial gradients in both liver and intestine. We further use isotope tracing to explore the spatial topography of metabolic activity.

These measurements were made using MALDI imaging mass spectrometry, an increasingly available tool^66^. Up to now, key limitations on MALDI metabolomics have been spatial resolution and signal-to-noise^67^. These are linked, as signal falls as the square of spatial resolution. At a spatial resolution of 15 µm in liver and 10 µm in intestine, we were able to decipher histological landmarks, classical gradients, and the single-cell layer of intestinal epithelium. This spatial resolution came at the expense of relatively low signal-to-noise and associated high pixel-to-pixel signal variation. Computationally, we deal with this lack of precision by a twist on averaging, using deep learning to discern, from data across roughly tens-to hundreds-of-thousands of MALDI pixels per image, the strongest topographical metabolic patterns. This enables robust detection of spatial patterns. Going forward, we foresee averaging across pixels remaining important, with the potential to do so in a manner that goes beyond metabolic gradients to pick up individual cell types, like Kupfer cells in the liver or goblet cells in intestine, likely empowered by multi-omic imaging^68–71^.

What shapes the observed spatial metabolic gradients? One simple explanation is that metabolite concentrations reflect enzyme expression. In the liver, we see alignment between enzyme expression and direct metabolic measurements, including for glutamine metabolism, TCA cycle feedstocks, and gluconeogenesis (Extended Data Fig. 3 and Extended Data Fig. 5). Focusing on glutamine, for example, glutamine catabolism to feed the urea and TCA cycles localizes periportally in terms of glutaminase transcript and enzyme (Extended Data Fig. 5c), glutamine flux contribution to TCA (Fig. 5b,c), and concentrations of the downstream metabolites glutamate and malate (Fig. 4a,b). In contrast, glutamine synthetase and its flux are pericentral (Fig. 5b and Extended Data Fig. 5c). In the intestine, while some metabolites follow associated enzymes (Extended Data Fig. 6e), others show striking opposing patterns (Extended Data Fig. 6a,b,f). For example, citrate is crypt localized, despite its synthetic enzyme citrate synthase being tip enriched and its consumption enzyme citrate lyase being crypt enriched (Extended Data Fig. 6a). Such discrepancies emphasize the importance of direct measurements of metabolite concentrations and flux.

A key reason why direct metabolic measurements are needed is that metabolism is shaped not just by enzymes, but also by substrate and oxygen availability^72^. As blood percolates from intestinal crypt to villus tip, and hepatic portal-to-central vein, oxygen is steadily consumed^46^. In the liver, both mitochondria and TCA intermediates colocalize in the periportal region with the more oxygen-rich blood. In the intestine, however, citrate localizes strongly to the oxygen-replete crypts, but mitochondria and other TCA intermediates localize to the relatively oxygen-poor tips. Notably, in both the liver and intestine, the mitochondria-rich areas surprisingly show lower energy charge, suggesting that these are areas of high, partially unsatiated energy demand.

As dietary nutrients pass from the intestinal lumen to systemic circulation, they are similarly steadily consumed. Through gavage of isotope-labeled fructose, we monitored this process with spatial resolution. Intestinal epithelium gets the first shot at dietary fructose. Rather than the geometrically privileged tips, we see faster fructose metabolism towards the bottom of villi. Intestinal epithelial cells process a portion of dietary fructose into glucose and organic acids. The remainder passes to the liver, where it first hits periportal cells. Both based on enzyme expression and our isotope labeling measurements, periportal liver is adept at catabolizing fructose, converting it to fructose-1-phosphate which rapidly breaks down into three-carbon units to drive energy production, gluconeogenesis, and glycogen storage. Thus, mammals seem to have evolved to clear fructose through a combination of intestinal and periportal metabolism, shielding the pericentral liver from most fructose.

For the large doses of fructose administered here, however, there is spillover of fructose to the pericentral liver, leading to accumulation of fructose-1-phosphate. This accumulation apparently arises due to active ketohexokinase but low aldolase B, the downstream enzyme. Such partial metabolism of fructose consumes ATP without regenerating it, leading to pericentral energy stress which is evident as local ATP depletion. Moreover, fructose-1 phosphate is a powerful lipogenic signal. This anatomical sequencing of fructose consumption has the benefit of rendering the fructose-1-phosphate-mediated lipogenic response insensitive to small quantities of fructose that are intestinally cleared (e.g. the first fruits of spring) and hypersensitive to large quantities that reach the pericentral liver (e.g. bountiful fruit in fall), meeting the evolutionary need to store fat selectively in times of sugar excess.

Overall, we provide foundational data regarding metabolite concentration gradients in the liver and small intestine. These are complemented by isotope tracer studies investigating glutamine, TCA cycle, and fructose metabolism. The ability to trace spatially the fates of dietary nutrients should be broadly applicable beyond fructose. More generally, the approaches presented here are poised for mapping metabolic gradients across a broad range of organs, including changes triggered by age and disease.

## Author contributions

This work was conceived by L.Z.S., B.J.R. and J.D.R. L.Z.S. developed the experimental MALDI-IMS methods and carried out most MALDI experiments. W.L helped with tissue washes. C.Z. and U.C. developed the deep-learning neural network model for mapping liver metabolic topography, with C.Z. innovating the multi-metabolic depth deep learning model. X.X and L.Z.S. developed supervised computational methods for portal-central axis construction. A.A. helped with intestinal MALDI-IMS and contributed to transcriptomics data analysis. Mouse experiments were done by L.Z.S., W.D.L., M.M., C.S.R.J. and C.J.H.

## Methods

### Animals

Mouse studies followed protocols approved by the Princeton University Animal Care and Use Committee. Animals were housed on a normal light cycle (8AM-8PM) and fed a standard rodent chow (PicoLab Rodent 20 5053, St. Louis, MO). Mice used were 12 – 18-week-old C57BL/6J males (The Jackson Laboratory). For spatial metabolomics experiments, mice were transferred to new cages with enrichment and water gel (ClearH2O, HydroGel) and without food at 8:30 AM, fasted for 7 hours and euthanized at 3:30 PM by cervical dislocation for tissue collection.

### Jugular vein catheterization

Aseptic surgery was done to insert a catheter (Instech Labs) in the right jugular vein which was connected to a vascular access button (Instech Labs) implanted under the back skin of the mouse. Catheterized mice were individually housed in environmentally enriched (Bed-r’Nest®, The Andersons) cages with ad libitum access to water and food. Mice were allowed to recover from jugular vein catheterization surgery for 5 days before experimentation. Catheters were flushed with heparin glycerol locking solution 5 days post-surgery.

### Glutamine infusion

[^13^C_5_-^15^N_2_]glutamine (99% ^13^C_5_, 99% ^15^N_2_, Cambridge Isotope Laboratories, CNLM-1275-H-0.5) solution was prepared in sterile saline at a concentration of 220 mM and filtered through a filter paper (pore size 0.45 µm). JV-catheterized mice were transferred to individual cages without food at 8 AM on the day of infusion. At 12 PM, mice were weighed to calculate the tracer infusion rate. Then, catheters were connected to infusion lines with a swivel and tether (Instech products: swivel SMCLA, line KVABM1T/25) that allowed the animals to move freely in the cage. The tracer solution was infused (SyringePump, NE-1000) at the specified rate for 5 min to fill the catheter dead volume 1 – 2 hours before starting the infusion, and mice were left to acclimatize. Infusions were started at 2 PM at a rate of 0.1 µl/min/g body weight for 2.5 h, and at 4:30 PM, animals were euthanized by cervical dislocation for tissue collection.

### Lactate infusion

[U-^13^C]L-Lactate tracer (98% ^13^C, 20 w/w% solution, CLM-1579, Cambridge Isotope Laboratories) was diluted to 5% in sterile water. Mice catheterized in the right jugular vein and left carotid artery were fasted by switching to a fresh cage with no food at 8 AM. At 12 PM, mice were weighed to calculate the tracer infusion rate, connected to the lines as above, and left in a cage for 1–2 h to acclimatize. At 14:30 – 15:30, infusion was initiated: a prime dose of 13 μl + 3 μl per gram mouse weight was provided over 22 s, and then the infusion rate was slowed to 0.3 μl min^−1^ per gram mouse weight. After 2.5 hours, mice were euthanized quickly by cervical dislocation, and tissues were collected for MALDI imaging.

### Fructose oral gavage

A 1:1 mixture of [D_12_]glucose (98%, DLM-9047-1, Cambridge Isotope Laboratories) and [U-^13^C]fructose (99%, CLM-1553-1, Cambridge Isotope Laboratories) in sterile saline was prepared. At 8:30 AM, mice were transferred to new cages without food. At 2:30 PM, mice were weighed to calculate the total volume to administer (10 μl/g body weight) and fed the tracer solution via a plastic feeding tube (Instech Laboratories, Plymouth Meeting, PA). Mice were euthanized by cervical dislocation either 90 seconds or 10 minutes post gavage and tissues were collected for MALDI imaging.

### Liver and small intestine collection for imaging mass spectrometry

For all liver imaging experiments, the left lateral liver lobe was collected within 1 minute after euthanasia, placed on a piece of aluminum foil, flash-frozen in liquid nitrogen and stored in a tightly sealed plastic bag at −80°C until workup for imaging.

Intestine collection and preparation for imaging mass spectrometry was optimized to preserve tissue integrity as follows: After animal euthanasia, the small intestine was collected by cutting 1 – 2 cm below the pyloric sphincter and taking 5 – 10 cm of the small-intestine length. The proximal end of the intestine was handheld and the tip of ∼1 mm-gauge stainless steel needle connected to a syringe prefilled with optimal cutting temperature embedding medium (OCT) (at room temperature) was gently inserted about 0.5 cm into the lumen. Space between the lumen and the needle tip was partially sealed by gently pressing with the fingers. OCT was pushed into the lumen until it filled about 1 cm of the intestine and the OCT-filled fragment was cut and placed into a 22 x 30mm x 20mm (width × length × depth) polyethylene disposable histology mold (Peel-A-Way® Tedpella) prefilled with ∼ 3 mm-high layer of OCT. Note that filling the intestine with OCT at high pressure (by pressing the syringe too hard) will result in tilted villi. Additional OCT was added to the mold to fully embed the intestines and placed on dry ice until it was fully frozen. The entire process took around 5 minutes. Finally, the tissue mold was stored in a tightly sealed plastic bag at −80 °C until processed.

### Tissue sectioning and preparation for MALDI-IMS

Frozen tissues were transferred from −80°C to a cryostat (Leica, CM3050S) at −20°C and left to thermally equilibrate for ∼30 min. For liver sectioning, frozen liver lobes were affixed by pouring a layer of OCT on cryostat stubs and placing the bottom of the lobe onto the OCT layer and left to freeze completely. For intestines, the OCT-embedded tissue block was removed from the mold and affixed to cryostat stubs as described for liver lobes.

Livers were sectioned in the transverse plane at 10-µm thickness and sections were thaw-mounted onto pre-cooled (to cryostat temperature) indium tin oxide (ITO)-coated slides. Intestine blocks were sliced either longitudinally (i.e., through the length of the intestine tubes) or across the intestine tubes (resulting in circular tissue section with the lumen at the center and villi peripheral) at 10 µm thickness and thaw-mounted onto pre-cooled ITO slides. Tissue sections on ITO slides were kept frozen on dry ice in preparation for MALDI matrix coating. Serial sections were collected and thaw-mounted on polylysine-coated glass slides and stored at −80°C for immunofluorescence.

### MALDI matrix coating

Right before matrix coating, tissue slices on ITO slides were dried under vacuum for 10 – 15 minutes. For liver MALDI imaging, we selected the MALDI matrix N-(1-Naphthyl) ethylenediamine dihydrochloride (NEDC) (product no. 222488, Sigma-Aldrich) and prepared a matrix solution at a concentration of 10 mg/ml in 70%:30% (v:v) methanol:water. Dried tissue sections were coated with NEDC using an automated HTX sprayer (model HTX or model M3+) (HTX Technologies, LLC). NEDC was sprayed at a flow rate of 0.1 ml/min, nozzle velocity of 1200 mm/min, nozzle temperature of 80°C, nozzle height of 40 mm, drying gas pressure of 10 psi, and a 30-s drying time between each pass, completing 10 passes over the entire slide.

For intestine MALDI imaging, we selected the matrix 1,5-DAN (1,5 diaminonaphthalene) (56451, Miilipore Sigma) at a concentration of 6 mg/ml in 40%:40%:20% (v:v:v) methanol:acetonitrile:water. Dried tissue sections were coated with DAN using an automated HTX sprayer (model M3+) (HTX Technologies, LLC) using the following spraying conditions: flow rate = 0.6 ml/min, nozzle velocity = 1200 mm/min, nozzle temperature = 75°C, nozzle height = 40 mm, drying gas pressure = 10 psi, and a 30-s drying time between each pass, for a total of 9 passes.

### MALDI-IMS

All MALDI imaging mass spectrometry experiments were done using a MALDI-2 timsTOF fleX (Bruker Daltonics GmbH & Co. KG) equipped with microGRID technology. Mass calibration was performed using a Tuning Mix solution (Agilent Technologies, G1969-85000) before starting the imaging run. For liver and intestine imaging, acquisitions were recorded in negative ion mode at a mass-to-charge ratio (*m/z*) range of 100 – 900. Energy difference between the collision cell and the quadrupole was set to 5 eV with a collision RF Vpp of 500 V. The quadrupole ion energy was set to 8 eV and the minimum m/z = 120, with the pre-TOF transfer time = 50 µs. Lock masses of *m/z* 124.0068 (taurine), *m/z* 255.2330 (palmitate), *m/z* 346.0558 (AMP), *m/z* 426.0222 (ADP), and *m/z* 514.2844 (taurocholic acid) were used for online calibration. Trapped ion mobility separation (TIMS) was not engaged during IMS runs. For all MALDI imaging experiments the laser Smart Beam was set to “Custom” mode while engaging “Beam scan”. For liver imaging, a single burst of 150 laser shots was delivered to each pixel at a frequency of 5 kHz in a pixel area of 15 × 15 µm^2^, and the raster step size was set to 15 µm in x and y directions. For imaging livers from fructose-gavaged and lactate- and glutamine-infused mice, spatial resolution was set to 20 µm and a total of 300 laser shots per pixel were used. For intestine imaging, a single burst of 75 shots were delivered at each pixel at a frequency of 5 kHz to a pixel area of 10 × 10 µm^2^ and the raster step size was set to 10 µm. For 5-µm and 6-µm spatial resolution imaging of intestines, 25 laser shots were used and the step size was set to 5 µm or 6 µm, respectively.

### Acidic methanol washing

Where signal enhancement for phosphates (e.g., ATP) was necessary in the small intestine, tissue sections were washed with acidic methanol following the protocol developed by Lu et al^53^. Briefly, hydrophobic barriers on both sides of the tissue section were drawn with a hydrophobic pen (H-4000, Vector Laboratories, Burlingame, CA, USA) and the slide was placed on an incline with cotton balls (Fisher 22-456-885) placed at the lower edge to adsorb the eluent. A total of 3 ml of ice-cold methanol with 0.05% (v:v) formic acid was pipetted onto the tissue sections and left to dry under vacuum before matrix coating.

### H&E staining post-MALDI-IMS

After MALDI-IMS analysis, the matrix was removed by placing the slide in ice-cold methanol for 5 minutes followed by 3 washes with 1xPBS for 5 minutes. Slides were briefly dipped in MilliQ water followed by immersion in hematoxylin for 10 minutes and then transferred to warm tap water for another 15 minutes. A second immersion in MilliQ water for 30 seconds was done and the slides were then dehydrated in a series of ethanol washes, beginning with 95% ethanol for 30 seconds, followed by eosin staining for 1 minute to highlight cytoplasmic components. After eosin application, slides were returned to 95% ethanol for 2 minutes, followed by 2 minutes immersion in 100% ethanol and another 2 minutes in xylene. Finally, the slide was sealed with Cytoseal 60 and a cover slip was applied. Slides were imaged using a BioTek Cytation 5 Cell Imaging Multimode Reader (Agilent, USA).

### Immunofluorescence staining

Liver tissue slices were fixed with 4% paraformaldehyde (PFA), washed with phosphate buffered saline (PBS) and incubated in PBS + 0.1% Triton-X for 1 - 2 hours for permeabilization. Tissue slices were then blocked with 0.1% bovine serum albumin (BSA) in PBS + 0.1% Triton-X containing for 1 hour and incubated with primary antibody (EpCAM/TROP1 antibody, BLR077G, NBP3-14685, Novus Biologicals, LLC) at 1:200 dilution for 2 hours at room temperature. Samples were then washed and incubated with fluorescent secondary antibodies (1:500 dilution) for 1 hour., followed by mounting with Fluoromount-G (Southern Biotech, 0100-01) and imaged using a BioTek Cytation 5 Cell Imaging Multimode Reader (Agilent, USA).

### MALDI-IMS data processing

Raw IMS data was converted to “.imZML” format using SCiLS^TM^ Lab (Bruker). The “.imZML” files were converted to “.mat” format using the in-house developed MATLAB software IsoScope (https://github.com/xxing9703/Isoscope)^32^. An initial set of roughly 800 – 1000 *m/z* peaks was extracted from the raw data using the untargeted peak picking feature in IsoScope. Briefly, this process filters peaks for which the mean signal across a subset of pixels is at least three times a predefined mean baseline. Since this process does not filter out background signal (including MALDI matrix peaks) and isotopologues, we matched *m/z* peaks with metabolites measured in-house by LC-MS analysis of liver and intestine extracts within a mass window of ± 15 ppm, yielding roughly 170 matching *m/z* peaks in liver and 100 in intestine. Of the 170 *m/z* peaks in liver, roughly 120 corresponded to deprotonated metabolites and lipids (the remaining roughly 50 peaks comprised the same compounds in different ion forms, e.g., [M+Na-2H]^-^ or were detectable in a minority of pixels). For the intestine, the list of about 100 *m/z* peaks corresponded to different deprotonated metabolites and lipids. A few ions measured only as chlorinated adducts or other ion forms were manually added to the lists. Ion intensities for the ∼170 spectral features in liver and the total ion count at each pixel were extracted and used for deep-learning analysis. Note that although glucose is one of several hexose isomers that cannot be distinguished in MALDI-TOF-IMS, glucose is the dominant isomer in the fasting liver based on liquid chromatography-mass spectrometry (LC-MS) (Extended Data Fig. 2a) (ratio of glucose to other hexoses > 100), and accordingly we refer to hexose in fasting liver as glucose.

### Deep learning of liver metabolic topography

We adapted GASTON^40^, an unsupervised deep learning algorithm originally developed for spatial transcriptomics, to spatial metabolomics data. GASTON learns a ‘topographic map’ of a 2-D tissue slice in terms of a 1-D coordinate called the *isodepth*, which is analogous to the height in a topographic map of a landscape. The key assumption of GASTON is that the expression of many genes is a function of the isodepth. Here, we introduce the related algorithm MET-MAP which learns a topographic map of a tissue slice from spatial metabolomics data by deriving a similar 1-D coordinate, which we call *metabolic depth*, under the assumption that the abundance of most metabolites is a function of the metabolic depth.

Briefly, spatial metabolomics data consists of a metabolite abundance matrix 𝐴 = [𝑎_𝑖,𝑚_] ∈ ℝ^𝑁×𝑀^ where entry 𝑎_𝑖,𝑚_ is the abundance of metabolite 𝑚 = 1, . . ., 𝑀 in spatial location 𝑖 = 1, . . ., 𝑁 and a spatial location matrix 𝑆 = [(𝑥_𝑖_, 𝑦_𝑖_)] ∈ ℝ^𝑁×2^ whose rows contain the 2D coordinates (𝑥_𝑖_, 𝑦_𝑖_) of location 𝑖 = 1, . . ., 𝑁. We aim to learn a continuously differentiable function 𝑑 ∶ ℝ^2^ → ℝ, which we call the *metabolic depth* function, and a continuously differentiable function ℎ_𝑚_ ∶ ℝ → ℝ for metabolite 𝑚𝑚, which we call the *1-D abundance function*, such that the observed abundance 𝑎_𝑖,𝑚_ is approximately equal to the composition ℎ_𝑚_(𝑑(𝑥_𝑖_, 𝑦_𝑖_)) of the metabolic depth function 𝑑𝑑 and the 1-D abundance function ℎ_𝑚_; that is 𝑎_𝑖,𝑚_ ≈ ℎ_𝑚_(𝑑(𝑥_𝑖_, 𝑦_𝑖_)). We do this by parameterizing the metabolic depth 𝑑 with a neural network and learning continuously differentiable functions ℎ_𝑚_ that minimize the mean squared error (MSE) between the observed abundance 𝑎_𝑖,𝑚_ and the predicted abundance ℎ_𝑚_(𝑑(𝑥_𝑖_, 𝑦_𝑖_)):

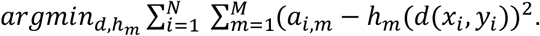

The value 𝑑(𝑥_𝑖_, 𝑦_𝑖_) gives the metabolic depth at spatial location (𝑥_𝑖_, 𝑦_𝑖_) and the gradient 𝛻𝑑(𝑥_𝑖_, 𝑦_𝑖_) indicates the spatial direction of maximum change in metabolite abundance. Details for the underlying model can be found elsewhere^40^.

Before training the neural network to solve for metabolic depth 𝑑, raw metabolite abundance [𝑎_𝑖,𝑚_] ∈ ℝ^𝑁×𝑀^ was first normalized using Total Ion Count (TIC) at individual pixels and log-transformed:

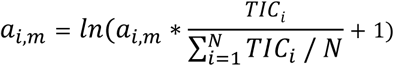

In liver samples, each tissue slice contained about 100,000 pixels. To ensure efficient training time and exclude low quality regions such as tissue tears and distorted veins, we manually divided each tissue slice into non-overlapping rectangular crops of size around 130 × 150 pixels. Each crop was then trained using the neural network to learn a metabolic depth for that specific crop. The metabolic depths of neighboring crops were well matched at the boundaries.

MET-MAP allows for different number of hidden layers and nodes within each layer to parameterize the functions 𝑑𝑑 ∶ ℝ^2^ → ℝ and ℎ_𝑚_ ∶ ℝ → ℝ . We obtained the lowest loss and efficient training times for metabolic depth function using a single hidden layer with 300-400 nodes, and the 1-D abundance functions using a hidden layer with 100 nodes. Since the metabolic depth is scale invariant, we standardized the scales by performing a few transformations for easier interpretation and analysis. First, the metabolic depth results were scaled so that the contour lines of metabolic depth were equally spaced on the physical space with a range of 0 to 1. Then, to ensure the metabolic depth following the direction of the portal-central axis such that the minimum depth corresponded to the portal node, we scaled it using bile acid (taurocholic acid), which is known to enrich consistently in the periportal regions. Specifically, we transformed metabolic depth 𝑑 to −𝑑 if *corr* (𝑑, taurocholic acid) < 0.

### Deep learning of intestinal epithelium metabolic topography

To isolate pixels that correspond to the epithelial layer in the intestine ion images, we performed k-means clustering (using the function in IsoScope) with input data of about 100 metabolites and lipids. This generated 4 clusters that corresponded to epithelial layer, lamina propria, lumen and background (mostly ITO slide and matrix). The epithelial cluster was isolated and ion intensities for those pixels passed to MET-MAP for deep learning of spatial gradients.

Unlike in the liver, initial manual inspection of raw metabolite abundance showed that metabolites in the intestine samples followed multiple distinct spatial patterns, two of the most dominant being the villus crypt-to-tip axis, and epithelial cell brush border-to-basal membrane axis (roughly bottom-to-top and outside-to-inside of villi, respectively). In order to model multiple distinct spatial patterns, we modified the neural network architecture of GASTON to learn 𝐾𝐾 metabolic depths instead of a single depth. Given metabolite abundances 𝐴 = [𝑎_𝑖,𝑚_] ∈ ℝ^𝑁×𝑀^, spatial coordinates 𝑆 = [(𝑥_𝑖_, 𝑦_𝑖_)] ∈ ℝ^𝑁×2^ and a parameter 𝐾, we learned a continuously differentiable function 𝑑: ℝ^2^ → ℝ^𝐾^, and *linear* functions ℎ_𝑚_: ℝ → ℝ, where each metabolite’s spatial pattern is best described by one of the 𝐾 metabolic depths 𝑑(𝑥_𝑖_, 𝑦_𝑖_) ∈ ℝ^𝐾^ learned. This soft assignment was achieved by making abundance mappings ℎ_𝑚_ linear and using lasso regularization to further sparsify the weights 𝑊 ∈ ℝ^𝑀×𝐾^ of the linear abundance mappings ℎ_𝑚_, where 𝑊_𝑘,𝑚_ represents the connection between the 𝑘-th metabolic depths and 𝑚𝑚-th metabolite abundance. Together, we learn:

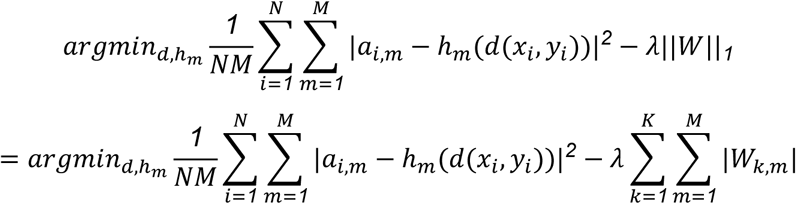

The 𝐾-dimensional latent variable 𝑑(𝑥_𝑖_, 𝑦_𝑖_) ∈ ℝ^𝐾^ models the 𝐾 distant spatial-metabolic patterns desired. We chose the value of the parameter 𝐾 using prior knowledge about the number of distinct patterns expected in the data. When the choice was unclear, we used 𝐾 = *4*. With effective lasso regularization on the weights connecting the 𝐾 metabolic depths with individual metabolites, we then excluded extra metabolic depths with close-to-zero weights that were not descriptive of metabolite abundance.

Similar to the data pre-processing in the liver, the intestine spatial metabolomics data was first log-transformed and normalized against TIC. The number of epithelial pixels in each intestine sample was similar to the number of pixels in each crop of the previous liver samples, so we did not further crop the intestine samples.

Based on the initial manual inspection of raw metabolite abundance, we set 𝐾𝐾 = *3* metabolic depths. Depending on individual sample quality as well as resolution, one or two of the metabolic depths were found to be significant with strong weights connecting to metabolite abundance. Through manual inspection, one of the significant metabolic depths was found to be the crypt-to-tip axis in every sample, validated by strong correlations with marker metabolites such as cholesterol sulfate and C20:3. The other significant metabolic depth, when present, correlated with the epithelial cell brush border-to-basal membrane axis, with strong correlation with metabolites such as linoleic acid. The metabolic depth results were then scaled similarly as in liver results, where minimum metabolic depth corresponded to crypt regions in the crypt-to-tip axis.

### Spatial gradient and statistics

After assigning metabolic depth values to pixels within a tissue crop, data was re-binned to 50 bins with equal number of pixels. Raw metabolite ion counts were normalized to the median count across all pixels within a single tissue crop. Then, normalized ion counts that correspond to the same metabolic depth bin were amalgamated from all crops across different mice. For liver, we excluded pixels that colocalize with portal and central veins (1 ≤ metabolic depth ≤ 7 for portal nodes and 48 ≤ metabolic depth ≤ 50 for central veins) to measure spatial gradients between the two nodes. Using the amalgamated liver data at 8 ≤ metabolic depth ≤ 47, normalized ion count as a function of metabolic depth was fitted to a linear model. Using predicted ion counts from the model at metabolic depth = 8 (periportal, PP) and 47 (pericentral, PC), we computed the fold change, PC/PP, which defines the porto-central gradient in liver. For intestines, we used predicted ion counts from linear regression at metabolic depth = 50 (tip) and 1 (crypt), and the fold change tip/crypt defines crypt-to-villus-tip gradient. *p*-values from linear regression were adjusted for false positives using the Benjamini-Hochberg correction procedure. For flux spatial gradients, ratios or fractions were calculated at each pixel classified by the neural network and downstream analysis was applied as described above.

### Ion count normalization and gradient plots

To plot metabolite intensity signal as a function of metabolic depth, pixels data was binned to 50 metabolic depth bins as described above. Within each tissue crop, for each metabolite, the median raw ion count at each bin was normalized to the overall median ion count for that metabolite in the tissue crop. Means were then taken across all tissue crops from the same mouse. Then, means and standard error of the mean were determined for each metabolite and bin across independent mice. Results were plotted for 8 ≤ metabolic depth ≤ 47 for liver and 1 ≤ metabolic depth ≤ 50 for intestine.

### Data processing for spatial isotope labeling

Data collected from mice infused with isotopically labeled lactate or glutamine or administered labeled fructose was converted to “.mat” as described above. Targeted metabolites of interest that were labeled by the tracer were added to the list of metabolites that were used in spatial metabolomics and passed to MET-MAP for analysis. After metabolic depth assignment, correction for natural abundance was first performed for each pixel using the software IsoScope as described previously^32^. Fractional labeling at each pixel was calculated as follows:

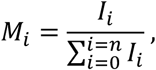

where M_i_ is the fraction of a given isotopologue, *n* is the number of carbon or nitrogen atoms in the metabolite of interest, and I_i_ is the raw ion count for a given isotopologue after natural isotope correction. Fractional labeling at each pixel was amalgamated across analyzed samples and analyzed by linear regression for spatial gradient measurement as described above.

### Image processing

Raw ion images from livers and root-mean-square-normalized (RMS) intestine images were exported from SCiLS^TM^ Lab (Bruker) as “OME-TIFF” files without selecting the “hotspot removal” feature and processed with ImageJ. The color scale was adjusted manually to optimize visualization. For liver, heme image was binarized by thresholding and subtracted from other metabolite images. The binarized heme image was used to create vein outlines and overlaid with the desired metabolite image. Vein outlines were colored olive-green for portal veins and wine-red for central veins based on manual inspection of the taurocholic acid and PI(38:4) signals, the former localizing adjacent to portal veins and the latter to central.

### Supervised porto-central axis construction

To validate the results from MET-MAP, we established an orthogonal computational approach to infer portal-central gradients based on supervised machine learning. This orthogonal approach for automated porto-central axis construction involves three main steps: vein identification, vein classification, and porto-central axis construction. Step 1 involves identifying central and portal veins, which is achieved by using ion images for heme, followed by 2D Gaussian filtering and binarization by thresholding. The positions of all vein centroids are registered. In step 2, we use an automated approach to classify the veins that were identified in step 1 as either central or portal. For each identified vein, a region of n × n pixel is cropped, with the vein being located at the center of the region. Ion intensities for heme, taurocholic acid, and arachidonic acid are extracted and the resulting 3-channel image is classified as portal or central by a pre-trained convolutional neural network model. This model uses high confidence, manually annotated veins as the training data and contains 6 layers: (a 3×3×3 convolution, a batchnorm, ReLU, 2 fully connected and a softmax layer). In step 3, all portal and central veins are connected by straight lines under bond length constraints (285 ≤ L ≤ 685 um).

### Blood collection and serum preparation

Tail blood was sampled from live mice before sacrificing, collected directly into Microvette-CB-300Z tubes (Sarstedt) and placed on ice. Tubes were centrifuged at 15,000 x g for 30 minutes at 4 °C, and serum was transferred to polypropylene tubes and stored at −80 °C.

Metabolites were extracted as previously described^6^. Briefly, 3-5 µL serum was added to 65 µL HPLC-grade methanol pre-cooled on wet ice, then vortexed and placed on dry ice for 15 minutes. The metabolite extract was then centrifuged at 15,000 x g, 15 minutes at 4 °C. The supernatant metabolite extract was diluted a further 5 to 10-fold in HPLC-grade methanol, centrifuged again, and the supernatant transferred to LC-MS tubes for analysis.

### Liquid-chromatography mass spectrometry

Metabolites were measured using a quadrupole-orbitrap mass spectrometer (QExactive Plus, Exploris 240, or Exploris 480, Thermo Fisher Scientific) operating in negative ion mode coupled to hydrophilic interaction liquid chromatography (HILIC) via electrospray ionization. Scans ranged from *m/z* 60 to 1000 at 1 Hz and a resolution ≥ 140,000. An XBridge BEH Amide column (2.1 mm × 150 mm, 2.5 µm particle size, 130 Å pore size; Waters) was used for LC separation with a gradient of solvent A (20 mM ammonium hydroxide in 95:5 water:acetonitrile, 20 mM ammonium acetate, pH 9.45) and solvent B (acetonitrile). Flow rate was set 150 µL/min. The LC gradient was: 0 min, 85% B; 2 min, 85% B, 3 min; 80% B; 5 min, 80% B; 6 min, 75% B, 7 min, 75% B, 8 min, 70% B; 9 min, 70% B, 10 min, 50% B; 12 min, 50% B; 13 min, 25% B; 16 min, 25% B; 18 min, 0% B, 23 min, 0% B, 24 min, 85% B. Autosampler temperature was 5 °C, and injection volume was set to 10 µL.

### LC-MS data processing

LC-MS data was analyzed using El-Maven (v.0.6.1). Ion counts were extracted and corrected for natural isotope abundance using the software isocorrCN (https://github.com/xxing9703/MIDview_isocorrCN). Serum enrichment was calculated as the ratio of the tracer abundance (i.e. infused isotopic form) to the sum of ion counts for all isotopologues after natural isotope correction.

## Notes

### Competing Interest Statement

The authors have declared no competing interest.

## References

1 Braeuning, A. et al. Differential gene expression in periportal and perivenous mouse hepatocytes. Febs Journal 273, 5051–5061 (2006). 10.1111/j.1742-4658.2006.05503.x

2 Halpern, K. B. et al. Single-cell spatial reconstruction reveals global division of labour in the mammalian liver. Nature 542, 352-+ (2017). 10.1038/nature21065

3 Jungermann, K. FUNCTIONAL-HETEROGENEITY OF PERIPORTAL AND PERIVENOUS HEPATOCYTES. Enzyme 35, 161–180 (1986). 10.1159/000469338

4 Mariadason, J. M. et al. Gene expression profiling of intestinal epithelial cell maturation along the crypt-villus axis. Gastroenterology 128, 1081–1088 (2005). 10.1053/j.gastro.2005.01.054

5 Moor, A. E. et al. Spatial Reconstruction of Single Enterocytes Uncovers Broad Zonation along the Intestinal Villus Axis. Cell 175, 1156-+ (2018). 10.1016/j.cell.2018.08.063

6 Hui, S. et al. Glucose feeds the TCA cycle via circulating lactate. Nature 551, 115-+ (2017). 10.1038/nature24057

7 Jang, C. et al. Metabolite Exchange between Mammalian Organs Quantified in Pigs. Cell Metabolism 30, 594-+ (2019). 10.1016/j.cmet.2019.06.002

8 Montoro-Huguet, M. A., Belloc, B. & Domínguez-Cajal, M. Small and Large Intestine (I): Malabsorption of Nutrients. Nutrients 13 (2021). 10.3390/nu13041254

9 Jang, C. et al. The Small Intestine Converts Dietary Fructose into Glucose and Organic Acids. Cell Metabolism 27, 351-+ (2018). 10.1016/j.cmet.2017.12.016

10 Gebhardt, R. METABOLIC ZONATION OF THE LIVER-REGULATION AND IMPLICATIONS FOR LIVER-FUNCTION. Pharmacology & Therapeutics 53, 275–354 (1992). 10.1016/0163-7258(92)90055-5

11 Trefts, E., Gannon, M. & Wasserman, D. H. The liver. Current Biology 27, R1147–R1151 (2017). 10.1016/j.cub.2017.09.019

12 Goodman, R. P., Calvo, S. E. & Mootha, V. K. Spatiotemporal compartmentalization of hepatic NADH and NADPH metabolism. Journal of Biological Chemistry 293, 7508–7516 (2018). 10.1074/jbc.TM117.000258

13 Katz, N. R. METHODS FOR THE STUDY OF LIVER-CELL HETEROGENEITY. Histochemical Journal 21, 517–529 (1989). 10.1007/bf01753352

14 Hoehme, S. et al. Prediction and validation of cell alignment along microvessels as order principle to restore tissue architecture in liver regeneration. Proceedings of the National Academy of Sciences of the United States of America 107, 10371–10376 (2010). 10.1073/pnas.0909374107

15 Jungermann, K. & Kietzmann, T. Zonation of parenchymal and nonparenchymal metabolism in liver. Annual Review of Nutrition 16, 179–203 (1996). 10.1146/annurev.nu.16.070196.001143

16 Harnik, Y. et al. A spatial expression atlas of the adult human proximal small intestine. Nature 632, 1101–1109 (2024). 10.1038/s41586-024-07793-3

17 Bahar Halpern, K., et al. Lgr5+telocytes are a signaling source at the intestinal villus tip. Nature Communications 11 (2020). 10.1038/s41467-020-15714-x

18 Allaire, J. M. et al. The Intestinal Epithelium: Central Coordinator of Mucosal Immunity. Trends in Immunology 39, 677–696 (2018). 10.1016/j.it.2018.04.002

19 Guerbette, T., Boudry, G. & Lan, A. Mitochondrial function in intestinal epithelium homeostasis and modulation in diet-induced obesity. Molecular Metabolism 63 (2022). 10.1016/j.molmet.2022.101546

20 Campbell, J., Berry, J. & Liang, Y. in Shackelford’s Surgery of the Alimentary Tract, 2 *Volume Set (Eighth Edition)* (ed Charles J. Yeo) 817–841 (Elsevier, 2019).

21 Ben-Moshe, S. et al. Spatial sorting enables comprehensive characterization of liver zonation. Nature Metabolism 1, 899–911 (2019). 10.1038/s42255-019-0109-9

22 Harnik, Y. et al. Spatial discordances between mRNAs and proteins in the intestinal epithelium. Nature Metabolism 3, 1680-+ (2021). 10.1038/s42255-021-00504-6

23 Hackett, S. R. et al. Systems-level analysis of mechanisms regulating yeast metabolic flux. Science 354 (2016). 10.1126/science.aaf2786

24 Stopka, S. A. et al. Spatially resolved characterization of tissue metabolic compartments in fasted and high-fat diet livers. Plos One 17 (2022). 10.1371/journal.pone.0261803

25 Seubnooch, P., et al. Characterisation of hepatic lipid signature distributed across the liver zonation using mass spectrometry imaging. Jhep Reports 5 (2023). 10.1016/j.jhepr.2023.100725

26 Tian, H. et al. Multimodal mass spectrometry imaging identifies cell-type-specific metabolic and lipidomic variation in the mammalian liver. Developmental Cell 59 (2024). 10.1016/j.devcel.2024.01.025

27 Jang, C., Chen, L. & Rabinowitz, J. D. Metabolomics and Isotope Tracing. Cell 173, 822–837 (2018). 10.1016/j.cell.2018.03.055

28 Bartman, C. R., TeSlaa, T. & Rabinowitz, J. D. Quantitative flux analysis in mammals. Nature Metabolism 3, 896–908 (2021). 10.1038/s42255-021-00419-2

29 Tran, D. H., et al. *De novo* and salvage purine synthesis pathways across tissues and tumors. Cell 187, 3602–3618.e3620 (2024). 10.1016/j.cell.2024.05.011

30 Pachnis, P. et al. In vivo isotope tracing reveals a requirement for the electron transport chain in glucose and glutamine metabolism by tumors. Science Advances 8, eabn9550 (2022). doi:10.1126/sciadv.abn9550

31 Yoon, S. J. et al. Comprehensive Metabolic Tracing Reveals the Origin and Catabolism of Cysteine in Mammalian Tissues and Tumors. Cancer Research 83, 1426–1442 (2023). 10.1158/0008-5472.Can-22-3000

32 Wang, L. et al. Spatially resolved isotope tracing reveals tissue metabolic activity. Nature Methods 19, 223-+ (2022). 10.1038/s41592-021-01378-y

33 Rabelink, T. J. et al. Analyzing Cell Type-Specific Dynamics of Metabolism in Kidney Repair. Journal of the American Society of Nephrology 33, 13–14 (2022).

34 Buglakova, E. et al. Spatial single-cell isotope tracing reveals heterogeneity of de novo fatty acid synthesis in cancer. Nature Metabolism 6, 1695–1711 (2024). 10.1038/s42255-024-01118-4

35 Wang, J. N. et al. MALDI-TOF MS Imaging of Metabolites with a N-(1-Naphthyl) Ethylenediamine Dihydrochloride Matrix and Its Application to Colorectal Cancer Liver Metastasis. Analytical Chemistry 87, 422–430 (2015). 10.1021/ac504294s

36 Dewez, F. et al. Precise co-registration of mass spectrometry imaging, histology, and laser microdissection-based omics. Analytical and Bioanalytical Chemistry 411, 5647–5653 (2019). 10.1007/s00216-019-01983-z

37 Liang, Z. L., Guo, Y. C., Sharma, A., McCurdy, C. R. & Prentice, B. M. Multimodal Image Fusion Workflow Incorporating MALDI Imaging Mass Spectrometry and Microscopy for the Study of Small Pharmaceutical Compounds. Analytical Chemistry 96, 11869–11880 (2024). 10.1021/acs.analchem.4c01553

38 Van de Plas, R., Yang, J. H., Spraggins, J. & Caprioli, R. M. Image fusion of mass spectrometry and microscopy: a multimodality paradigm for molecular tissue mapping. Nature Methods 12, 366–U138 (2015). 10.1038/nmeth.3296

39 Molenaar, M. R. et al. Increasing quantitation in spatial single-cell metabolomics by using fluorescence as ground truth. Frontiers in Molecular Biosciences 9 (2022). 10.3389/fmolb.2022.1021889

40 Chitra, U. et al. Mapping the topography of spatial gene expression with interpretable deep learning. bioRxiv, 2023.2010.2010.561757 (2023). 10.1101/2023.10.10.561757

41 Yang, G. Y. et al. Glucuronidation: driving factors and their impact on glucuronide disposition. Drug Metabolism Reviews 49, 105–138 (2017). 10.1080/03602532.2017.1293682

42 Allain, E. P., Rouleau, M., Lévesque, E. & Guillemette, C. Emerging roles for UDP-glucuronosyltransferases in drug resistance and cancer progression. British Journal of Cancer 122, 1277–1287 (2020). 10.1038/s41416-019-0722-0

43 Wang, Y. F. & Chen, H. R. Protein glycosylation alterations in hepatocellular carcinoma: function and clinical implications. Oncogene 42, 1970–1979 (2023). 10.1038/s41388-023-02702-w

44 Blomme, B., Van Steenkiste, C., Callewaert, N. & Van Vlierberghe, H. Alteration of protein glycosylation in liver diseases. Journal of Hepatology 50, 592–603 (2009). 10.1016/j.jhep.2008.12.010

45 Ljungqvist, O. Human Metabolism; A Regulatory Perspective. Clinical Nutrition 39, 3531–3531 (2020). 10.1016/j.clnu.2020.09.019

46 Kietzmann, T. Metabolic zonation of the liver: The oxygen gradient revisited. Redox Biology 11, 622–630 (2017). 10.1016/j.redox.2017.01.012

47 Manco, R. & Itzkovitz, S. Liver zonation. Journal of Hepatology 74, 466–468 (2021). 10.1016/j.jhep.2020.09.003

48 Kater, J. M. Comparative and experimental studies on the cytology of the liver. Zeitschrift für Zellforschung und Mikroskopische Anatomie 17, 217–246 (1933). 10.1007/BF00374042

49 Kang, S. W. S. et al. A spatial map of hepatic mitochondria uncovers functional heterogeneity shaped by nutrient-sensing signaling. Nature Communications 15 (2024). 10.1038/s41467-024-45751-9

50 Angerer, T. B., Bour, J., Biagi, J.-L., Moskovets, E. & Frache, G. Evaluation of 6 MALDI-Matrices for 10 μm Lipid Imaging and On-Tissue MSn with AP-MALDI-Orbitrap. Journal of the American Society for Mass Spectrometry 33, 760–771 (2022). 10.1021/jasms.1c00327

51 Baliou, S. et al. Protective role of taurine against oxidative stress (Review). Mol Med Rep 24, 605 (2021). 10.3892/mmr.2021.12242

52 Strott, C. A. & Higashi, Y. Cholesterol sulfate in human physiology: what’s it all about? Journal of Lipid Research 44, 1268–1278 (2003). 10.1194/jlr.R300005-JLR200

53 Lu, W. Y. et al. Acidic Methanol Treatment Facilitates Matrix-Assisted Laser Desorption Ionization-Mass Spectrometry Imaging of Energy Metabolism. Analytical Chemistry 95, 14879–14888 (2023). 10.1021/acs.analchem.3c01875

54 Johnson, R. J. et al. Potential role of sugar (fructose) in the epidemic of hypertension, obesity and the metabolic syndrome, diabetes, kidney disease, and cardiovascular diseased. American Journal of Clinical Nutrition 86, 899–906 (2007).

55 Abdelmalek, M. F. et al. Increased Fructose Consumption Is Associated with Fibrosis Severity in Patients with Nonalcoholic Fatty Liver Disease. Hepatology 51, 1961–1971 (2010). 10.1002/hep.23535

56 Chung, M. et al. Fructose, high-fructose corn syrup, sucrose, and nonalcoholic fatty liver disease or indexes of liver health: a systematic review and meta-analysis. American Journal of Clinical Nutrition 100, 833–849 (2014). 10.3945/ajcn.114.086314

57 Sievenpiper, J. L. et al. Effect of Fructose on Body Weight in Controlled Feeding Trials A Systematic Review and Meta-analysis. Annals of Internal Medicine 156, 291–U291 (2012). 10.7326/0003-4819-156-4-201202210-00007

58 Tsilas, C. S. et al. Relation of total sugars, fructose and sucrose with incident type 2 diabetes: a systematic review and meta-analysis of prospective cohort studies. Canadian Medical Association Journal 189, E711–E720 (2017). 10.1503/cmaj.160706

59 Jensen, T. et al. Fructose and sugar: A major mediator of non-alcoholic fatty liver disease. Journal of Hepatology 68, 1063–1075 (2018). 10.1016/j.jhep.2018.01.019

60 Bartman, C. R. et al. Slow TCA flux and ATP production in primary solid tumours but not metastases. Nature 614, 349–357 (2023). 10.1038/s41586-022-05661-6

61 Jang, C. et al. The small intestine shields the liver from fructose-induced steatosis. Nature Metabolism 2, 586-+ (2020). 10.1038/s42255-020-0222-9

62 Heinz, F., Lamprecht, W. & Kirsch, J. ENZYMES OF FRUCTOSE METABOLISM IN HUMAN LIVER. Journal of Clinical Investigation 47, 1826-+ (1968). 10.1172/jci105872

63 Adelman, R. C., Ballard, F. J. & Weinhouse, S. Purification and Properties of Rat Liver Fructokinase. Journal of Biological Chemistry 242, 3360–3365 (1967). 10.1016/S0021-9258(18)95917-X

64 Kagimoto, T. & Uyeda, K. Hormone-stimulated phosphorylation of liver phosphofructokinase in vivo. Journal of Biological Chemistry 254, 5584–5587 (1979). 10.1016/S0021-9258(18)50449-X

65 Underwood, A. & Newsholme, E. PROPERTIES OF PHOSPHOFRUCTOKINASE FROM RAT LIVER AND THEIR RELATION TO THE CONTROL OF GLYCOLYSIS AND GLUCONEOGENESIS. Biochemical Journal 95, 868–875 (1965). 10.1042/bj0950868

66 Alexandrov, T. Spatial metabolomics: from a niche field towards a driver of innovation. Nature Metabolism 5, 1443–1445 (2023). 10.1038/s42255-023-00881-0

67 Buchberger, A. R., DeLaney, K., Johnson, J. & Li, L. Mass Spectrometry Imaging: A Review of Emerging Advancements and Future Insights. Analytical Chemistry 90, 240–265 (2018). 10.1021/acs.analchem.7b04733

68 Hu, T. et al. Single-cell spatial metabolomics with cell-type specific protein profiling for tissue systems biology. Nature Communications 14, 8260 (2023). 10.1038/s41467-023-43917-5

69 Esselman, A. B. et al. In situ molecular profiles of glomerular cells by integrated imaging mass spectrometry and multiplexed immunofluorescence microscopy. Kidney International (2024). 10.1016/j.kint.2024.11.008

70 Nunes, J. B. et al. Integration of mass cytometry and mass spectrometry imaging for spatially resolved single-cell metabolic profiling. Nature Methods 21, 1796–1800 (2024). 10.1038/s41592-024-02392-6

71 Vicari, M. et al. Spatial multimodal analysis of transcriptomes and metabolomes in tissues. Nature Biotechnology 42, 1046–1050 (2024). 10.1038/s41587-023-01937-y

72 Vander Heiden, M. G., Cantley, L. C. & Thompson, C. B. Understanding the Warburg Effect: The Metabolic Requirements of Cell Proliferation. Science 324, 1029–1033 (2009). doi:10.1126/science.1160809

73 Thomsson, K. A. et al. Sulfation of O-glycans on Mucin-type Proteins From Serous Ovarian Epithelial Tumors. Molecular & Cellular Proteomics 20, 100150 (2021). 10.1016/j.mcpro.2021.100150

