## Extended Data Figures for "Spatial metabolic gradients in the liver and small intestine"

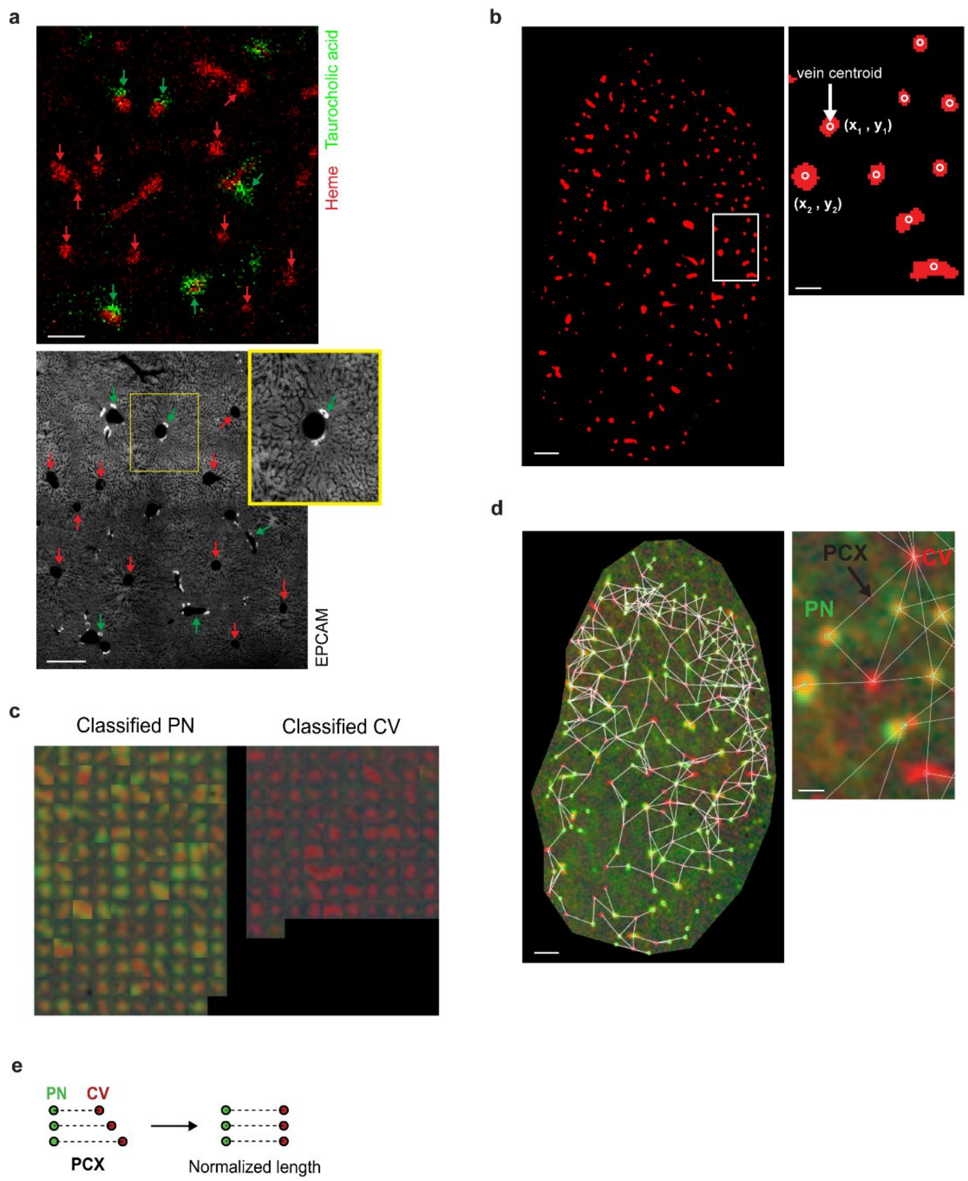


**Extended Data Figure 1. Supervised construction of portal-central axes based on MALDI-IMS.**

**(a)** Top: Overlaid MALDI images of heme (red) and taurocholic acid (green). Bottom: Immunofluorescence image of EPCAM in a serial section. EPCAM is primarily expressed on cells lining bile ducts near portal veins (green arrow in inset). Red arrow: central vein, green arrow: portal node. Scale bar = 200 μm (top), 300 μm (bottom)

**(b)** Central and portal veins are mapped using heme image (see Methods). Veins are registered and their centroids are assigned. Scale bar = 600 μm; inset scale bar = 150 μm.

**(c)** A subset of vein crops displaying signal for heme (red), taurocholic acid (green) and C20:4 (shown faintly in blue) is used to train a convolutional neural network (CNN), with the model output classifying each vein as central or portal. The same model is then used to classify all registered veins across the entire tissue slice.

**(d)** Classified portal and central veins are connected by portal-central axes under bond length constraints. Scale bar = 600 μm; inset scale bar = 150 μm.

**(e)** Physical lengths of the connecting lines are normalized to merge data across all axes.

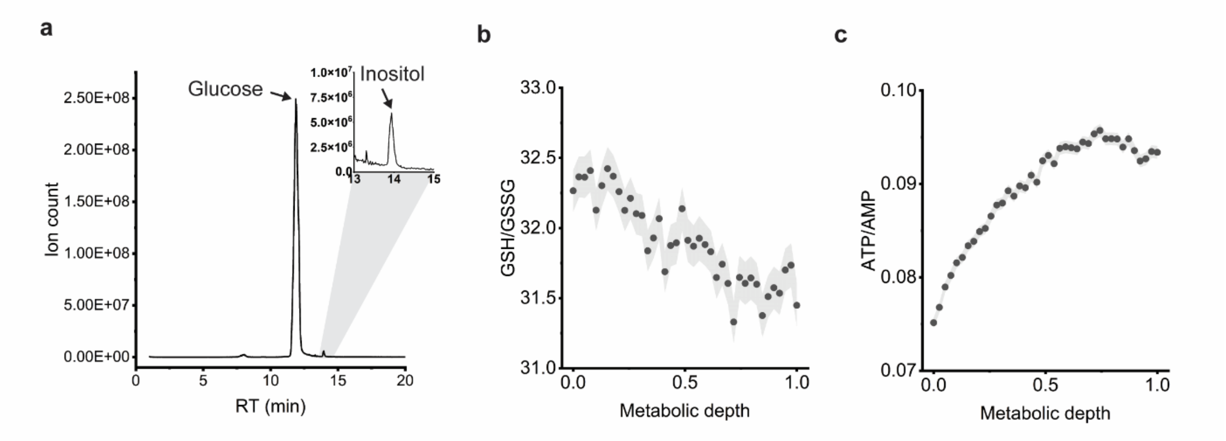


**Extended Data Figure 2. Glutathione-to-oxidized-glutathione (GSH/GSSG) and ATP-to-AMP ratios show opposite spatial gradients in the liver.**
**(a)** Glucose is the dominant hexose in the liver as measured by LC-MS of liver extracts (from fasted mice).
**(b)** GSH/GSSG ratio shows periportal localization in the liver ($p$ = 0 from linear regression, data merged across n = 7 independent mice).
**(c)** Same as in (b) but for ATP/AMP ratio ($p$ = 0).

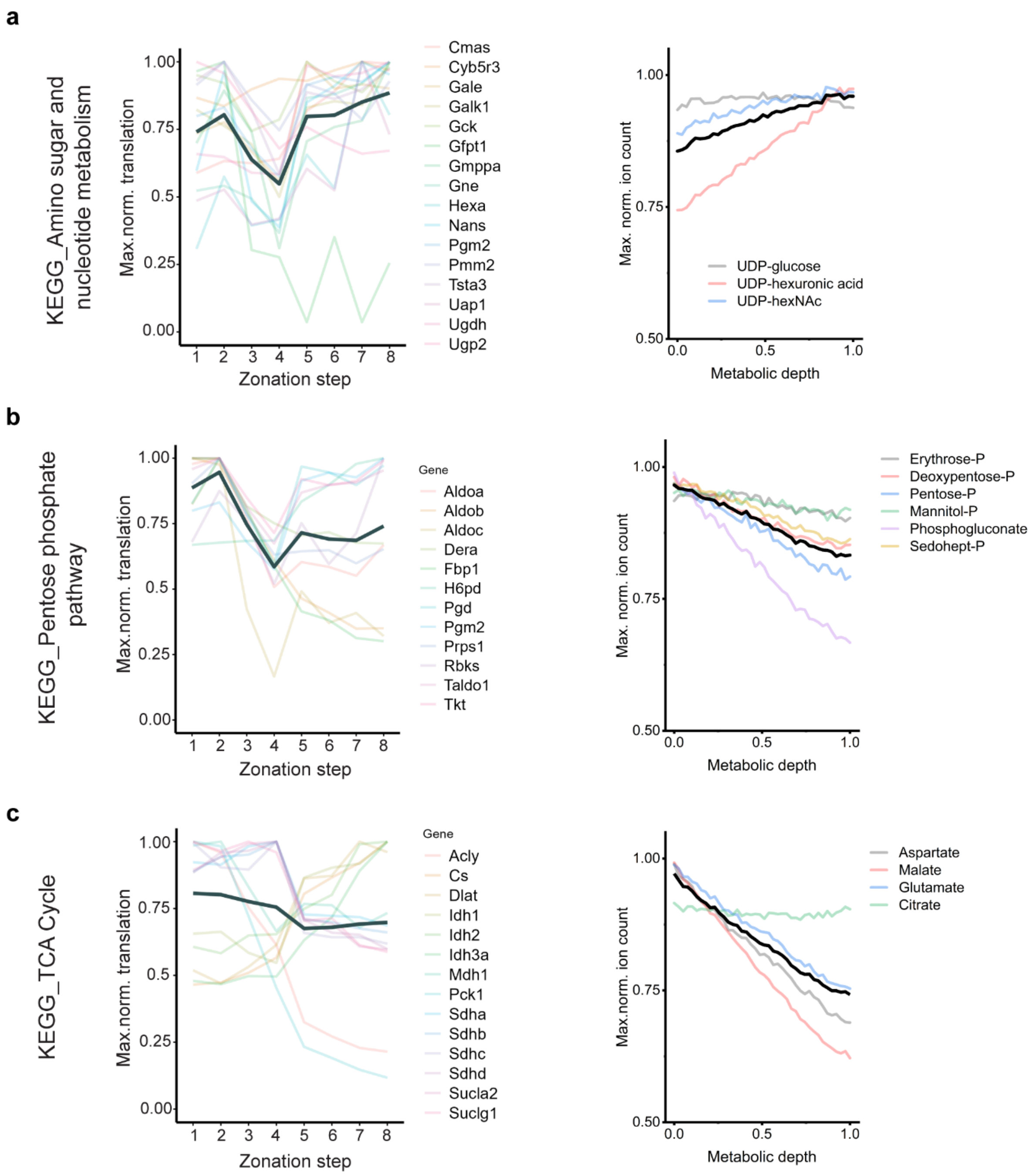


**Extended Data Figure 3. Spatial correlation between protein and metabolite abundances in liver.**
**(a)** Portal-central zonation profiles for proteins involved in amino sugar and nucleotide metabolism (left) and associated metabolites measured by MALDI-IMS (right). Zonation steps from 1 – 8 (left) represent portal-to-central direction. Each protein is represented by a color as annotated by the corresponding gene name (right panel). Data for both proteins and metabolites were normalized to maximum abundance. Black bold lines represent mean values across shown proteins or metabolites. Protein data published by Ben-Moshe et al[21].
**(b)** Same as in (a) for pentose phosphate pathway.
**(c)** Same as in (a) for TCA cycle.

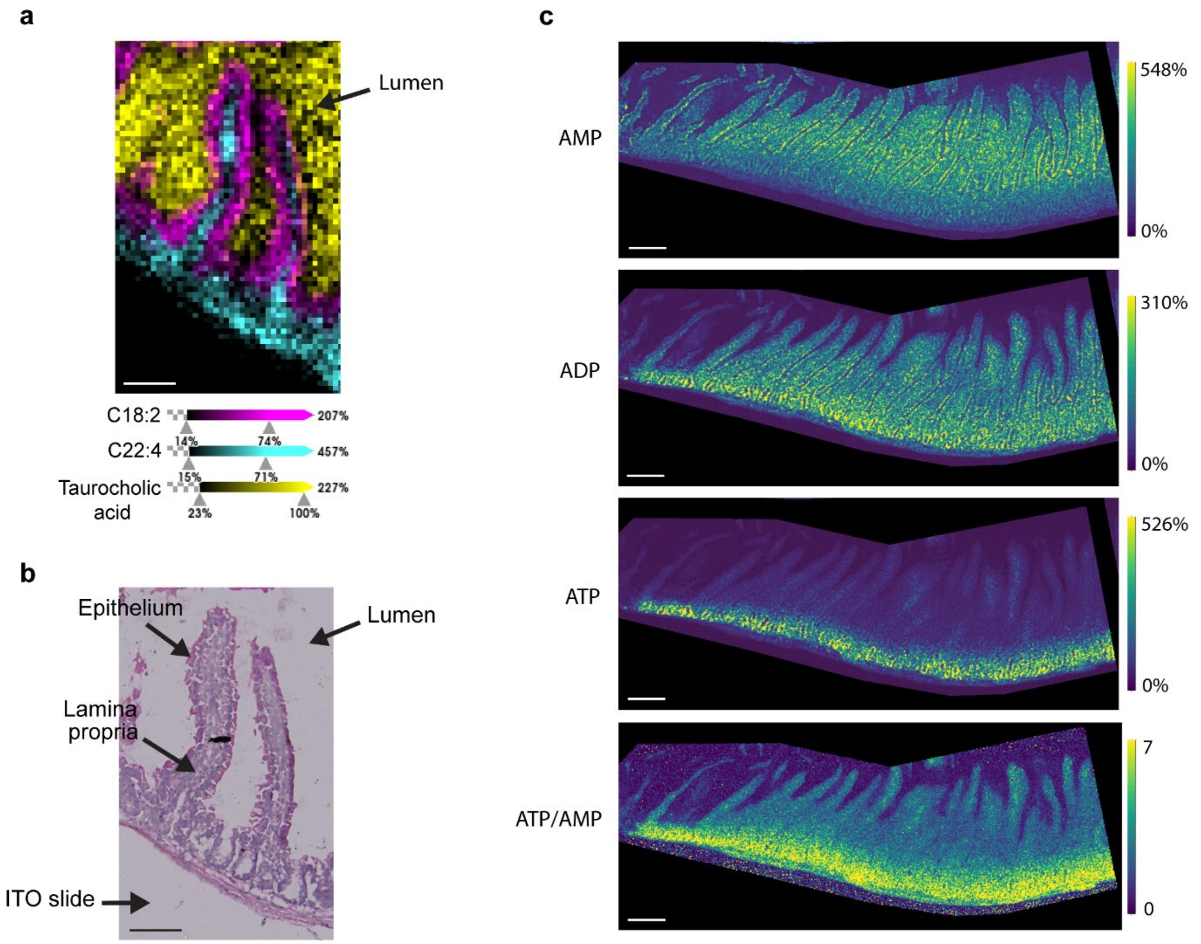


**Extended Data Figure 4. Intestinal crypts display higher ATP-to-AMP ratio.**
**(a)** Overlaid raw MALDI images of C18:2 (magenta) marking the epithelial layer, C22:4 (cyan) marking the lamina propria, and taurocholic acid (yellow) marking the lumen.
**(b)** H&E of the same intestine tissue section as in (a) after MALDI-IMS.
**(c)** AMP (top), ADP (second from top) and ATP (third from top) raw images in intestines and ATP/AMP ratio (bottom) after tissue section was washed with methanol + 0.05% formic acid. MALDI image in (a) collected at 10×10 $\mu m^2$ and (c) 5×5 $\mu m^2$ spatial resolution. Scale bars: (a – b) 100 μm; (c) 200 μm.

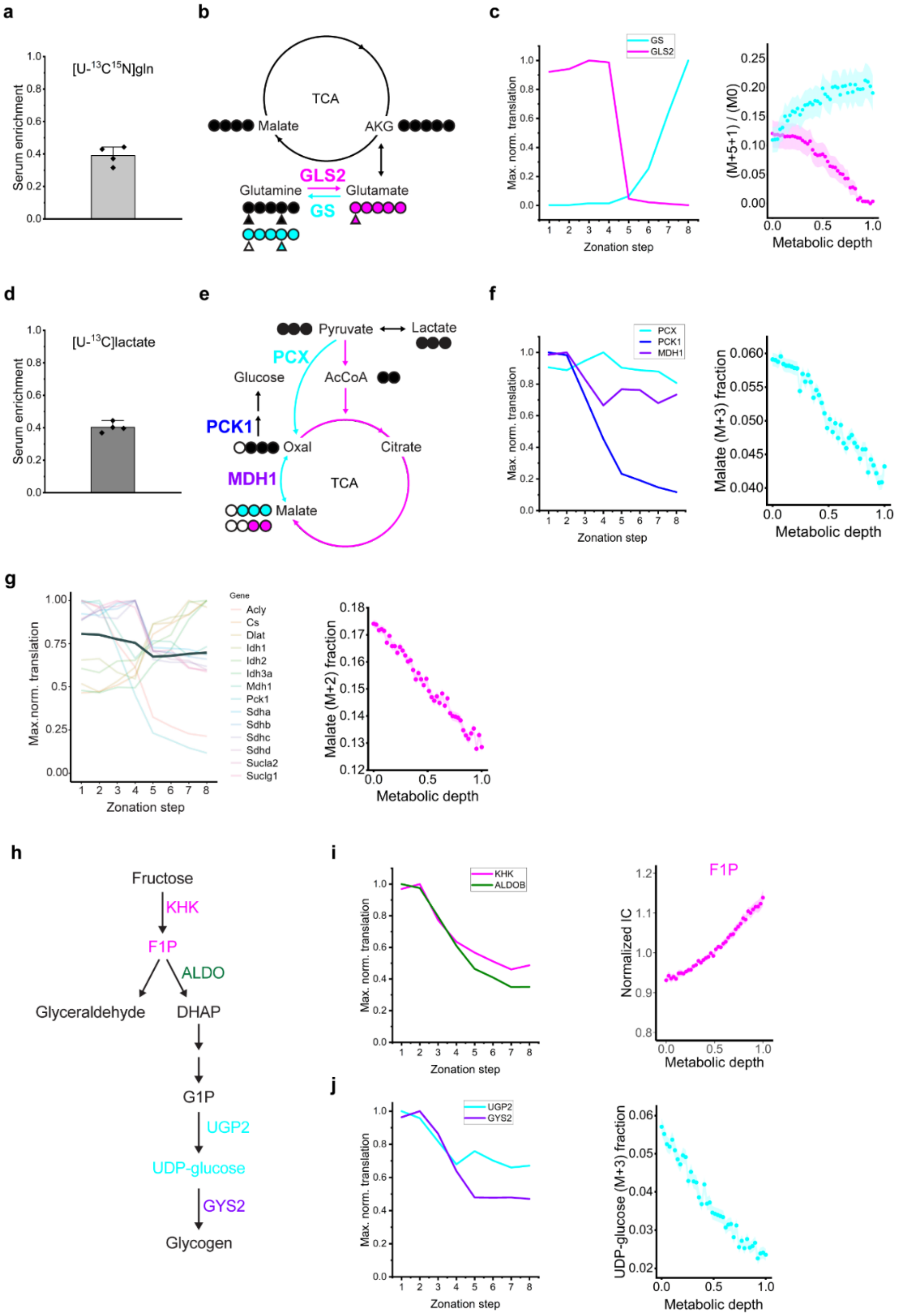


**Extended Data Figure 5. Spatial correlation between protein abundances and fluxes in liver.**

**(a)** [U-$^{13}$C,$^{15}$N]glutamine serum enrichment from JV-infusions of n = 4 independent mice.

**(b)** Schematic of [$^{13}C_5$,$^{15}N_2$]glutamine tracer fates: catabolism to glutamate by GLS2 (magenta) and recycling glutamine via glutamine synthetase (cyan). Circles: carbon atoms. Triangles: nitrogen atoms. Filled (both colored and black): isotope-labeled.

**(c)** Portal-central zonation profiles for the enzymes GLS2 and GS (left) and gradients of the associated glutamine and glutamate labeling patterns (right). Data same as in Figure 5b.
**(d)** [U-$^{13}$C]lactate serum enrichment from JV-infusions of n = 4 independent mice.
**(e)** Schematic of TCA-cycle labeling from [U-$^{13}$C]lactate. Lactate oxidation generates M+2 malate through TCA turning. Anaplerosis produces M+3 malate.
**(f)** Zonation profiles of analplerotic enzymes (left) and gradient of malate M+3 fraction (right). Data same as in Figure 5f.
**(g)** Zonation profiles of TCA-cycle enzymes (left) and gradient of malate M+2 fraction (right). Data same as in Figure 5e.
**(h)** Schematic of dietary fructose utilization in the liver.
**(i)** Zonation profiles of KHK and ALDOB (left) and gradient of F1P (right). Data same as in Figure 6c.
**(j)** Zonation profiles of glycogenic enzymes (left) and gradient of UDP-glucose (M+3) fraction (right) produced from carbon-labeled oral fructose. Data same as in Figure 6c. All protein data shown published by Ben-Moshe et al[21].

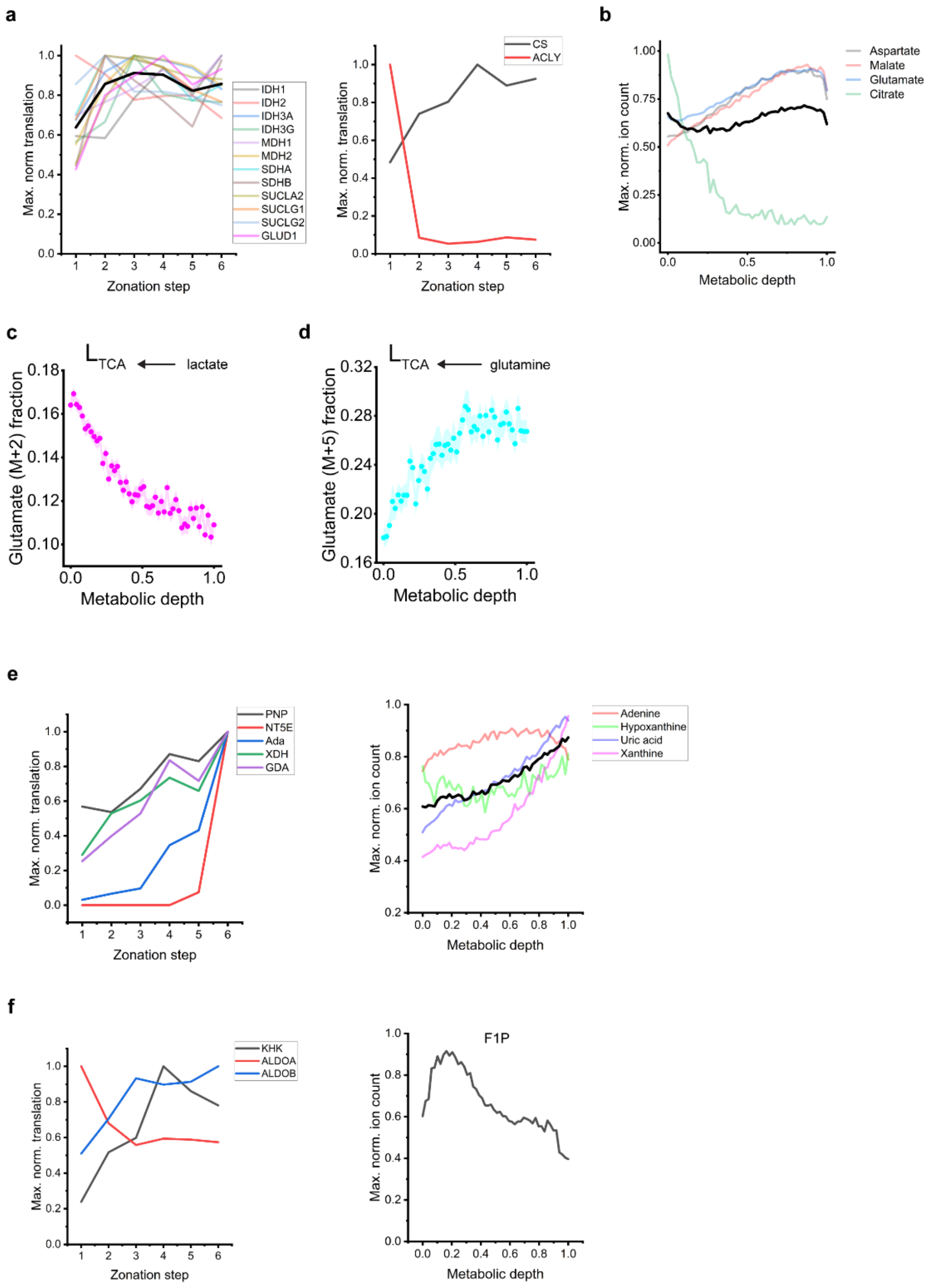


**Extended Data Figure 6. Spatial correlations between protein, metabolite abundances and fluxes in intestine.**

**(a)** TCA-cycle enzymes are higher in tip than crypt. Black bold line shows mean across plotted enzymes. The citrate-producing enzyme, citrate synthase (CS), is enriched in the tips, whereas the citrate-consuming enzyme, ATP citrate lyase (ACLY), is enriched in the villus bottom (right panel).

**(b)** Most TCA-cycle intermediates are higher in tip than crypt, with the exception of citrate. Black bold line shows mean across plotted metabolites.

**(c)** Circulating lactate feeds TCA preferentially in crypts. Data same as in Figure 5h.

**(d)** Circulating glutamine feeds TCA preferentially in tips. Data same as in Figure 5j.

**(e)** Purine degradation enzymes localize in villus tips (left panel). Purine degradation products localize in the villus tips (right panel). Black bold line shows mean of plotted metabolites. Data same as in Figure 4a.
**(f)** The two enzymes involved in fructose metabolism, KHK and ALDOB, localize in tips (left panel), whereas F1P localizes in villus bottom (right panel). Data same as in Figure 6b. Protein data are from Harnik et al[22].

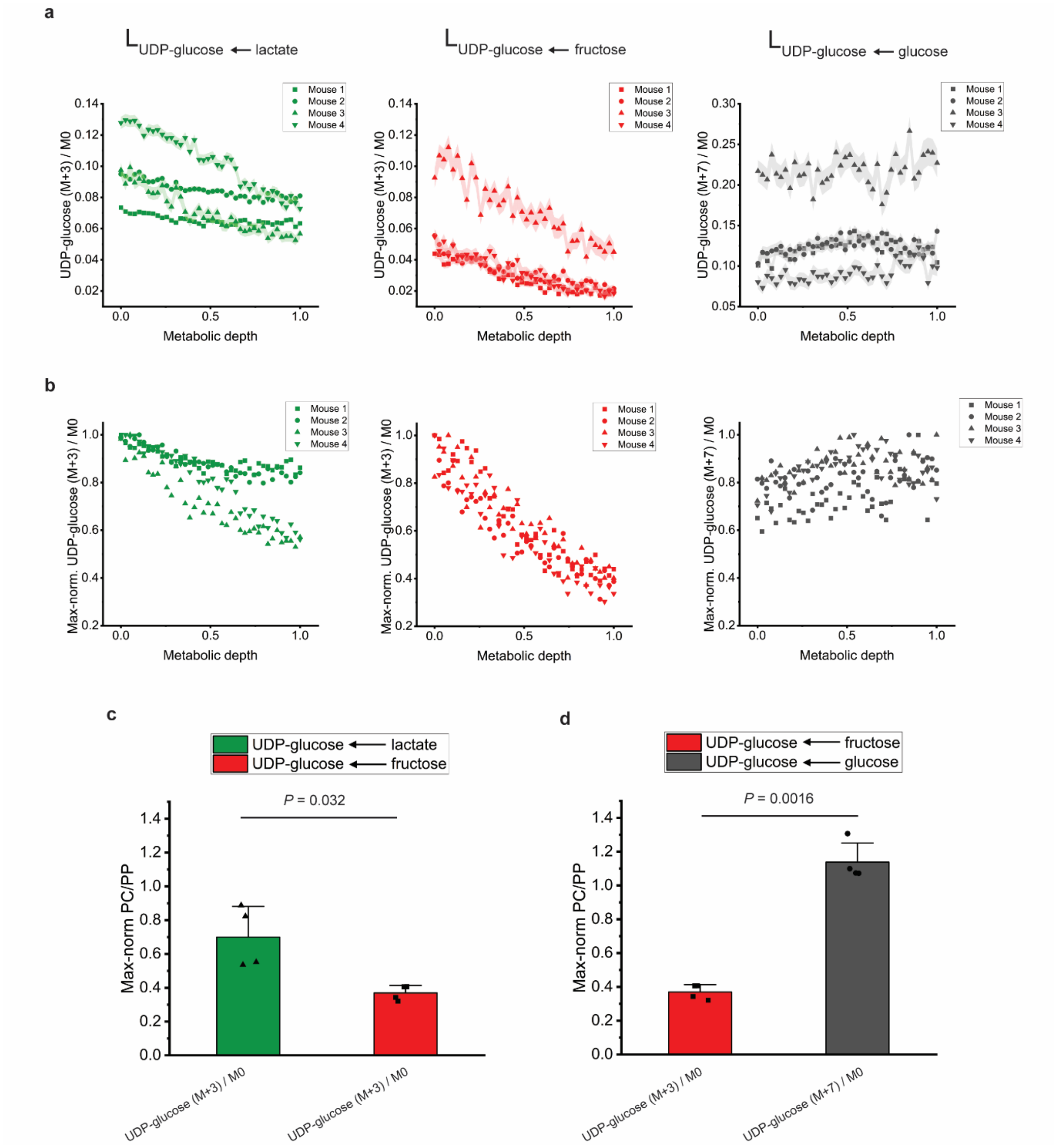


**Extended Data Figure 7. UDP-glucose production from oral labeled fructose is periportal.**
**(a)** Ratios of labeled to unlabeled UDP-glucose in liver from IV infusion of $^{13}$C-lactate (left), and oral $^{13}$C-fructose (middle) and oral $^{13}$C-glucose (right). Each symbol represents mean ratio from one mouse (n = 4 mice).
**(b)** Normalized data from (a), replotted after normalizing to the maximum labeling.
**(c)** Data in (b) is fitted to a linear model and the ratio of slopes from the regression, reported as pericentral (PC) to periportal (PP) values, are compared by two-sample t-test (two-tailed).
**(d)** Same as (c) but comparing UDP-glucose labeling from oral-fructose (red) and co-administered oral glucose (gray).
